# Machine learning-based clustering of nanosized fluorescent extracellular vesicles

**DOI:** 10.1101/2020.11.27.374728

**Authors:** Sören Kuypers, Nick Smisdom, Isabel Pintelon, Jean-Pierre Timmermans, Marcel Ameloot, Luc Michiels, Jelle Hendrix, Baharak Hosseinkhani

## Abstract

Extracellular vesicles (EV) are biological nanoparticles that play an important role in cell-to-cell communication. The phenotypic profile of EV populations is a promising reporter of disease, with direct clinical diagnostic relevance. Yet, robust methods for quantifying the biomarker content of EV have been critically lacking, and require a single-particle approach due to their inherent heterogeneous nature. Here, we used multicolor single-molecule burst analysis microscopy to detect multiple biomarkers present on single EV. We classified the recorded signals and applied the machine learning-based t-distributed stochastic neighbor embedding algorithm to cluster the resulting multidimensional data. As a proof of principle, we applied the method to assess both the purity and the inflammatory status of EV, and compared cell culture and plasma-derived EV isolated via different purification methods. We then applied this methodology to identify intercellular adhesion molecule-1 (ICAM-1) specific EV subgroups released by inflamed endothelial cells, and to prove that apolipoprotein-a1 is an excellent marker to identify the typical lipoprotein contamination in plasma. Our methodology can be widely applied on standard confocal microscopes, thereby allowing both standardized quality assessment of patient plasma EV preparations, and diagnostic profiling of multiple EV biomarkers in health and disease.

---

Extracellular vesicles (EV), biological nanometer-sized lipid bilayer enveloped particles, are intercellular communicators carrying specific biomarkers such as lipids, proteins and nucleic acids.^1,2^ Owing to their unique nature, EV are progressively becoming key players in health and disease. EV are both promising nanosized carriers in therapy but are equally important as diagnostic and/or prognostic tools.^3^ Blood plasma is a rich source of biomarkers and identifying disease-associated EV biomarkers in plasma has been challenging due to the presence of co-isolates such as protein complexes or lipoproteins.^4^ Moreover, the release of EV by almost every cell type into body fluids leads to a rich variety of biomarkers carrying EV subsets, making the characterization and downstream analysis of disease-specific EV biomarkers challenging.^5^ To assess these problems, the different biological nanoparticle populations need to be differentiated to enable (1) a purity check-up of EV fractions, as defined by the minimal information for studies of extracellular vesicles (MISEV) guidelines^6^ and (2) the phenotyping of multiple biomarkers on single EV, both crucial strategies to reliably identify EV-specific subgroups and/or biomarker panels. The MISEV guidelines define a series of control experiments to be executed for the reliable characterization of EV preparations. In these experiments, the size distribution and concentration of the anoparticles in the isolated EV samples is quantified. Furthermore, the presence of conventional membrane-bound markers *e.g*. the tetraspanins CD9, CD63and CD81 and intraluminal EV markers *e.g*. Annexin II as well as the absence of possible coisolates *e.g*. non-EV related proteins or lipoproteins are analyzed.

Conventional methodologies applied in diagnostics are not suited for phenotyping and characterizing single nanometer sized EV due to requirements imposed by the detection limit and spatial resolution of common instruments. To overcome these problems, the EV field has been intensively searching for translational methodologies in recent years.^7^

A powerful high-resolution technology is single burst analysis spectroscopy (SBA).^8^ In SBA picomolar fluorescently labeled nanoparticles diffuse freely and one-at-a-time through the small focal (femtoliter) volume on a confocal microscope. This in turn generates time-dependent fluorescence traces containing light bursts that are recorded. These single bursts can be identified and analyzed, which provides information about the respective single nanoparticles. Wyss e*t al*. first employed SBA to study the size profile and CD63 protein expression by using fluorescein isothiocyanate (FITC) labeled anti-CD63 antibody based immunostaining.^9^ However, this can be improved to fully exploit the potential of SBA. The use SBA with multiple markers will enable the identification of biological nanoparticle populations within the EV sample preparations.

The machine learning algorithm t-distributed stochastic neighbor embedding (t-SNE), a non-linear dimensionality reduction tool, allows the capture of local relationships between a set of data points and is often used in single-cell RNA sequencing data to identify specific cell populations.^10,11^ t-SNE is also applied in high-dimensionality fluorescence activated cell sorting (FACS) experiments, where this method can be used for automated gating or to identify different cell populations.^12^ The ability to capture patterns in multidimensional data could allow the identification of (rare) EV subpopulations.

In this study, we perform multicolor SBA to detect the colocalization of multiple fluorescently labeled markers on the surface of a single EV and use machine learning to find patterns to discern EV subgroups. We first characterized our EV preparations according to the MISEV guidelines. Next, we demonstrated the use of SBA on fluorescent nanoparticles and afterwards on fluorescently labeled EV. Recorded EV signals were classified and clustered using t-SNE. We have proven that our proposed SBA methodology can reliable determine the purity of plasma EV isolated comparable to the labor intensive and sample consuming conventional methods. Moreover, we applied the methodology to identify and quantify an inflammation associated EV subpopulation of diagnostic relevance.

## Results

### Preparation and standard (MISEV) characterization of cell- and plasma-derived EV (cEV and pEV)

In order to check the presence, purity and quality of EV preparations, a series of standard quality control experiments were performed on both plasma-derived EV (pEV) and cell-derived EV (cEV) fractions according to the MISEV 2018 guidelines (figure 1).^6^ Accordingly, the purity of the EV fractions was confirmed using western blotting assays for a classical EV marker (CD9), an EV cytosolic marker (Annexin II) and a non-EV marker (APOa1 for pEV and Golgin subfamily A member 2 (GM130) for cEV) as negative control (figure 1A). EV size distribution, concentration (figure 1B, E and H) and morphology (figure 1C, F and I) were evaluated using nanoparticle tracking analysis (NTA) and transmission electron microscopy (TEM), respectively. NTA uses light scattered by nanoparticles that diffuse by Brownian motion to identify sample size distributions. TEM allows us to study the single particle nature of EV. NTA demonstrated a mode of 116 ±1 nm for the single step isolation procedure of pEV (Figure 1B), a mode of 95 ±3 nm for the two-step isolation procedure of pEV (figure 1E) and a mode of 106 ±7 nm for cEV (figure 1H). As reported before, western blot analysis revealed the presence of APOa1 (28 kDa) positive lipoproteins along with typical EV markers (CD9, 25 kDa and Annexin II, 37 kDa) in the SEC-based isolated pEV (figure 1A), while the APOa1 marker was not detected in the pEV fraction (fractions 9 and 10) after applying a two-step procedure (SEC + Optiprep™ density gradient (ODG)) (figure 1D).^5^ In case of cEV isolates, the purity of samples was confirmed by the absence of the non-EV marker as a negative control (GM130, 130 kDa) in the SEC fraction (figure 1G). Using TEM, we confirmed the presence of cup shaped vesicles in both pEV and cEV fractions (figure 1C-F-I). In addition to the western blot results, TEM images also showed less aggregates after applying a two-step isolation procedure for EV from plasma (figure 1F). Overviews for these samples are provided in the supplemental information (SI figure 1). Together, these results confirmed that pure fractions of EV were isolated from both citrated plasma and cell conditioned medium according to the MISEV guidelines.

**Figure 1.**
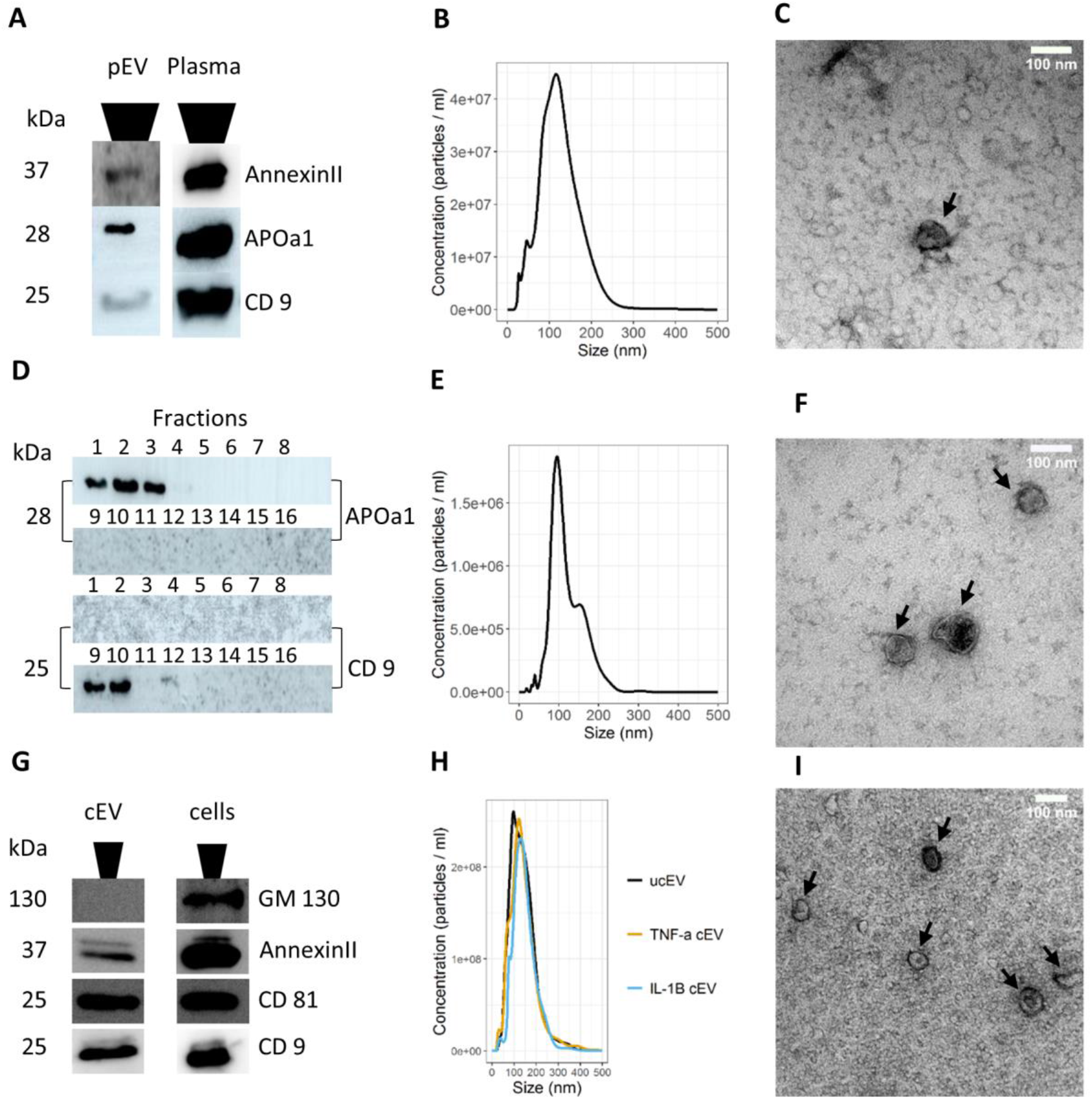
Quality control of EV isolates from citrated plasma (pEV) and cell conditioned medium (cEV) using a single step size exclusion chromatography (SEC) for pEV and cEV or Optiprep™ density gradient (ODG) combined with SEC for pEV according to the MISEV guidelines. **A**. Western blot analysis for classic EV markers CD9 (25 kDa), annexin II (37 kDa) and APOa1 (28 kDa) showing the presence in the pEV sample. Lysed citrated plasma is used as a positive control. **B**. NTA shows a distinguished peak between 100 and 200 nm, corresponding to the size of EV. **C**. Transmission electron microscopy (TEM) micrograph of plasma SEC isolated fractions, revealing pEV (black arrow) and co-isolates (scale bar 100 nm). **D**. Western blot analysis of the different one ml ODG column fractions. APOa1 (28 kDa) is enriched in the first three fractions (APOa1(+)-pEV fractions) and the EV marker CD9 (25 kDa) is enriched in the EV fractions 9 and 10 (CD9(+)-pEV fractions). **E**. NTA of the SEC isolated pEV from ODG fractions 9 and 10, demonstrates a distinguished peak around 100 nm and a shoulder around 175 nm. **F**. TEM micrograph of ODG isolated fractions, visualizing clearly distinguishable pEV (black arrows), with a higher purity (scale bar 100 nm). **G**. Western blot analysis showing the presence of annexin II (37 kDa), CD 81 (25 kDa) in cEV (SEC), as well as the absence of GM130 (130 kDa) a non-EV marker. **H**. NTA sizing results of cEV coming from treated human umbilical vein endothelial cells (HUVEC) (tumor necrosis factor alpha (TNF α) cEV and interleukin 1 beta (IL-1β) cEV) as well as coming from untreated HUVEC (ucEV), demonstrates a size range between 100 and 300 nm consistent with EV size. TEM micrograph of SEC isolated fractions, also showing clearly distinguishable cEV (black arrows, scale bar 100 nm).

### SBA detects multiple fluorescently labeled 100 nm beads

To evaluate the conceptual potential and sensitivity of SBA for measuring the size of small particles (∼100 nm), we employed commercially available FITC labeled silica beads.

With SBA, single bursts were identified and classified. This principle is further illustrated in figure 2. First, we use NTA to confirm the size of particles. This revealed that the mean size of the FITC labeled silica beads was approximately 138 ±1 nm and a mode of 128 ±15 nm (figure 3A). The histogram of the burst duration, the time a single nanoparticle needs to diffuse through the focal volume, provides information about the average size of the nanoparticles in the sample (figure 2B). SBA resulted in one major peak with a mean size of 111 ±1 nm and a mode of 104 ±1 nm (figure 3B, gray bars). These obtained results are similar to the NTA results, confirming that SBA can be used to obtain size distribution data for nanoparticles. Next, we analyzed commercially available multicolor TetraSpeck™ beads using SBA to assess the capability of detecting more than one label on a single nanoparticle. TetraSpeck™ beads are fluorescent nanoparticles containing four well-separated excitation/emission peaks (360/430 nm, 505/515 nm, 560/580 nm and 660/680 nm). Again, the size of these TetraSpeck™ beads was measured using NTA, giving a mean size of 118 ±5 nm, a mode of 83 ±2 nm. The SBA determined mean size was 125 ±3 nm and a mode of 96 ±14 nm (figure 3B, red bars).We then compared the different fluorescence signals of the TetraSpeck™ beads in two channels using two channel detection. As shown in figure 3C, we compared the beads (in black) with a single labeled sample (in red). This demonstrated that the signals of the single labeled sample were solely observed in one channel, whereas the signals of the individual TetraSpeck™ beads were-divided between the two channels.

**Figure 2.**
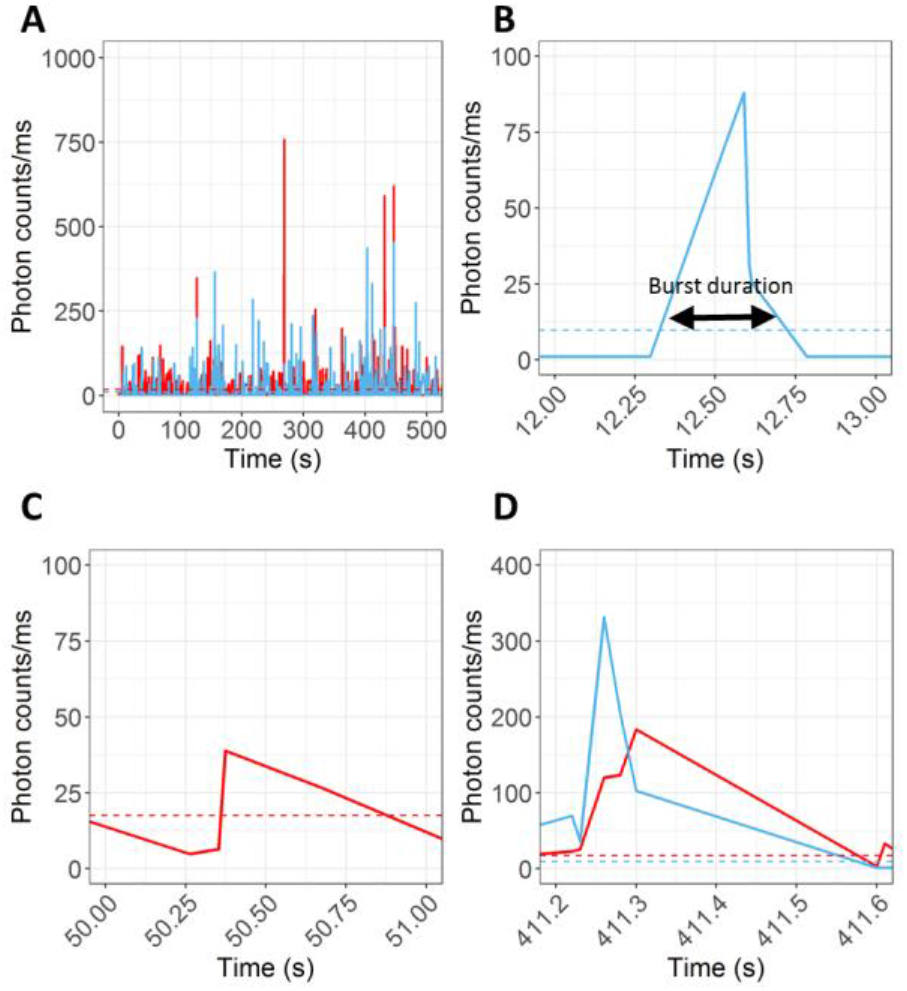
Graphical representation of SBA showing the SBA analysis principle a representative plasma derived EV sample is analyzed, with in red the fluctuations measured in the 488 nm channel and in blue the fluctuations measured in the 543 nm channel. A. Visualizes a typical time trace observed for a sample in two channels, a red channel and a blue channel. The calculated threshold is indicated by the red and blue dashed line. Any burst above the threshold is positive for that specific label(s). B. Visualization of a single burst in the blue channel. Everything below the dashed red line will be taken as noise and omitted from analysis. The black arrow represents the burst duration. C. Visualization of a single burst in the red channel. D. Visualization of a dual burst in both the red and the blue channel.

**Figure 3.**
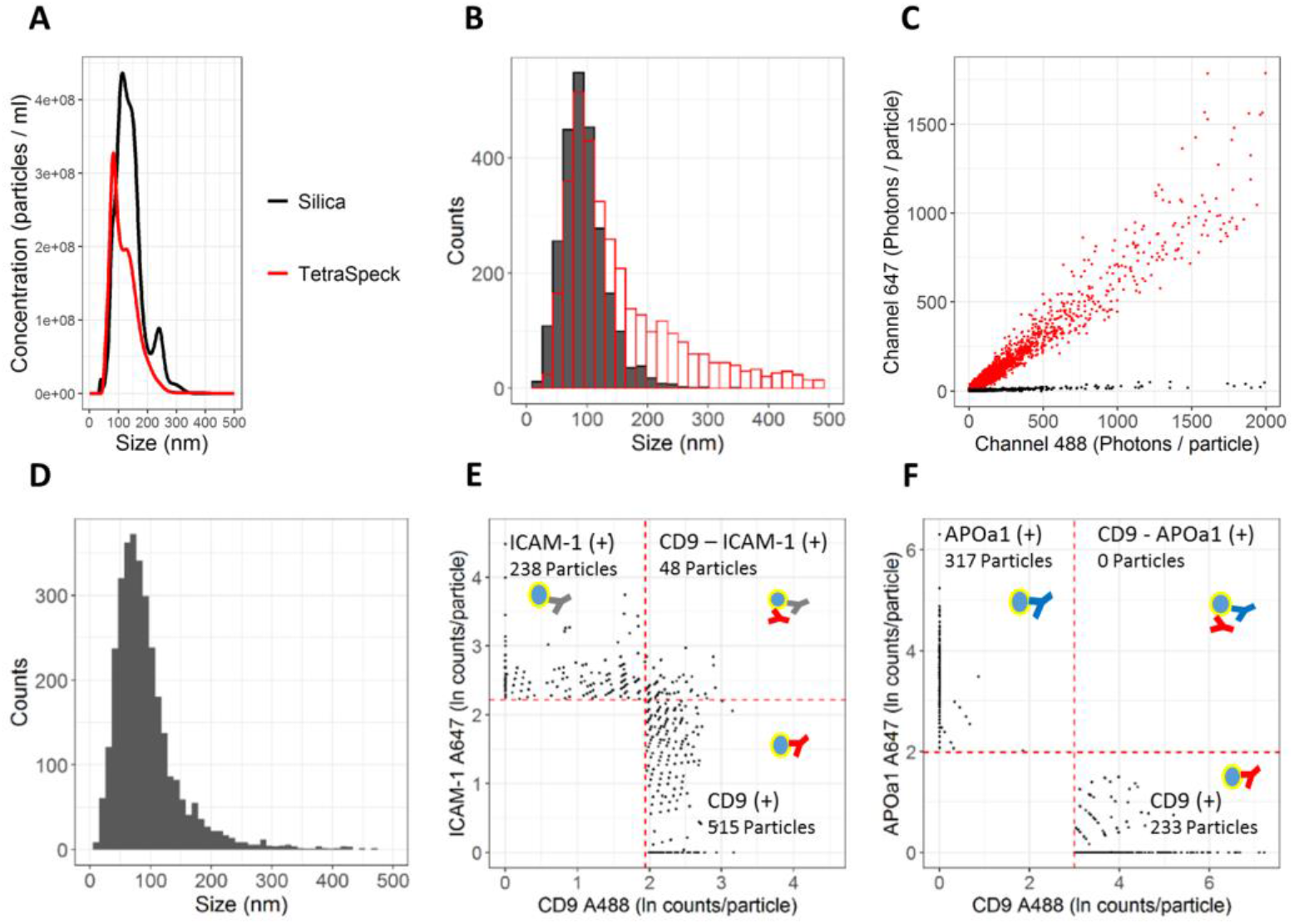
NTA and SBA based sizing and SBA profiling of beads and dual colored pEV preparations. A. NTA measurement of 100 nm FITC labeled silica beads (black) and TetraSpeck™ beads (red) both demonstrating a peak around 100 nm, as well as a second peak between 200 and 300 nm. B. SBA size distribution for 100 nm silica beads (black) and TetraSpeck™ beads (red) both visualizing a peak around 100 nm. C. SBA measurement of 100 nm TetraSpeck™ beads (red) and a single labeled sample (black) in two channels, clearly differentiating between both samples and enabling the visualization of TetraSpeck™ beads detected in two channels. D. SBA size plot of pEV Alexa 488 labeled with anti-CD9 (CD9 A488) and Alexa 647 labeled anti-ICAM-1 (ICAM-1 A647) visualizes a peak around 100 nm. E. Scatterplot of the sample in E. Thresholds for a positive burst are indicated in red, enabling the identification of single EV (CD9(+) or ICAM-1(+)) and dual positive EV (CD9 and ICAM-1(+)). F. Scatterplot of pEV labeled with CD9 A488 and Alexa 647 labeled anti-APOa1 (APOa1 A647) showing distinct CD9(+)- EV and the APOa1(+)-lipoproteins

These results confirmed that SBA provides information on the size distribution of particles. Furthermore, it can be applied to identify multiple labels on the same single bead.

### SBA allows the simultaneous detection of two or more surface markers on single EV and other biological nanoparticles

After demonstrating the applicability of the SBA approach on beads, we further investigated whether SBA can be used to detect different (surface) proteins on a single EV. To this extent, we first labeled EV with a combination of two different fluorescently labeled antibodies: Alexa 488 labeled anti-CD9 and Alexa 647 labeled anti-ICAM-1 (figure 3E) or Alexa 488 labeled anti-CD9 and Alexa 647 labeled anti-APOa1 (figure 3F). As shown in figure 3D, the SBA determined EV size distribution displayed a mean size of 95 ±2 nm corresponding with a mean burst duration of 2 ± 0.1 ms for pEV dually labeled with anti-CD9 Alexa 488 and anti-ICAM-1 Alexa 647 antibodies. Additionally, we independently confirmed that this duration is related to the diffusion constant, and hence to the hydrodynamic radius of the EV using fluorescence correlation spectroscopy (SI figure 2). Then, we adjusted the signals in each detection channel for spectral overlap of the used fluorophores and then defined the threshold for the detection of bursts in each individual channel (*e.g*. for Alexa 488 and Alexa 647) as described in the materials and methods and the number of detected photons in each channel was plotted. The thresholds allowed to divide the plot into quadrants and hence to identify individual nanoparticles as either single or double labeled. This is demonstrated by labeling pEV with labeled antibodies for CD9 and ICAM-1. As presented in figure 3E, a larger part of pEV carried only a single marker including ICAM-1 (26.6 ±2.2%) and CD9 (70.2 ±3.6%), while only 3.2 ±1.6% of pEV contained both ICAM-1 and CD9 markers. These results show that SBA can detect multiple surface markers on a single EV.

In relation to the quality of our EV preparations, our western blot analysis showed that the APOa1 marker (a key candidate marker of lipoproteins and potential co-isolate in EV preparations) is still present in the pEV isolated using one-step procedure (figure 1A). This sample was also analyzed using SBA. Based on the SBA, 57 ±3% and 43 ±3% of pEV also contained APOa1 and CD9 markers, respectively (figure 3F). No double labeled EV were detected, indicating that no APOa1 and CD9 labeled antibodies were simultaneously present on the detected nanoparticles. This demonstrates that with the proposed SBA approach two separate biological groups of nanoparticles within one sample can easily be identified, in this case lipoproteins and EV.

### t-SNE determined clusters correspond to EV subpopulations

To further explore the potential of SBA-based EV subpopulation profiling, we next evaluated whether our SBA-based approach can detect three markers on a single EV. To this end, pEV labeled anti-CD9, phycoerythrin (PE) labeled anti-CD63 and Alexa 647 labeled anti-ICAM-1 antibodies. Samples were prepared either using a mixture of these antibodies or with each of these antibodies separately with a subsequent pooling step, SI figure 3 visualizes the obtained bursts from BurstBrowser (PAM).

First SBA signals were classified and quantified as a percentage of the total number of detected bursts, thereby comparing the separately labeled with the triple labeled (multiplex) pEV sample, as shown in figure 4A. The quantified single labeled EV are comparable for both the separately labeled and the triple labeled (multiplex) samples. The multiplex labeled sample shows percentages between 1 and 6% for two and three labels, whereas the separately labeled sample has dual-labeled percentages below 0.75%. To visualize the potential of EV subpopulation detection within the data, t-SNE was applied. This machine learning algorithm enables the identification of clusters, *i.e*. potential EV subpopulations within multidimensional data. Combining the t-SNE clusters with the classification data allows the visualization of different EV subpopulations. Figure 4B gives a simplified intuitive example on how with t-SNE clusters of the original high dimensional dataset are preserved in a lower dimension. Figure 4C shows the t-SNE analysis of the separately labeled sample, resulting in three major clusters of EV profiles. However, in these single-labeled clusters some EV were indicated as double- or triple-labeled EV, most probably as a result of the applied least squares unmixing method. In contrast, figure 4D shows the results of the triple labeled (multiplex) sample clearly demonstrating the presence of dual- or triple-labeled EV. The overviews are also given in the supporting information (SI figure 4). More information on the optimization of these clusters is provided in SI figure 5. As expected, no clusters can be identified for the TetraSpeck™ beads (SI figure 6).

**Figure 4.**
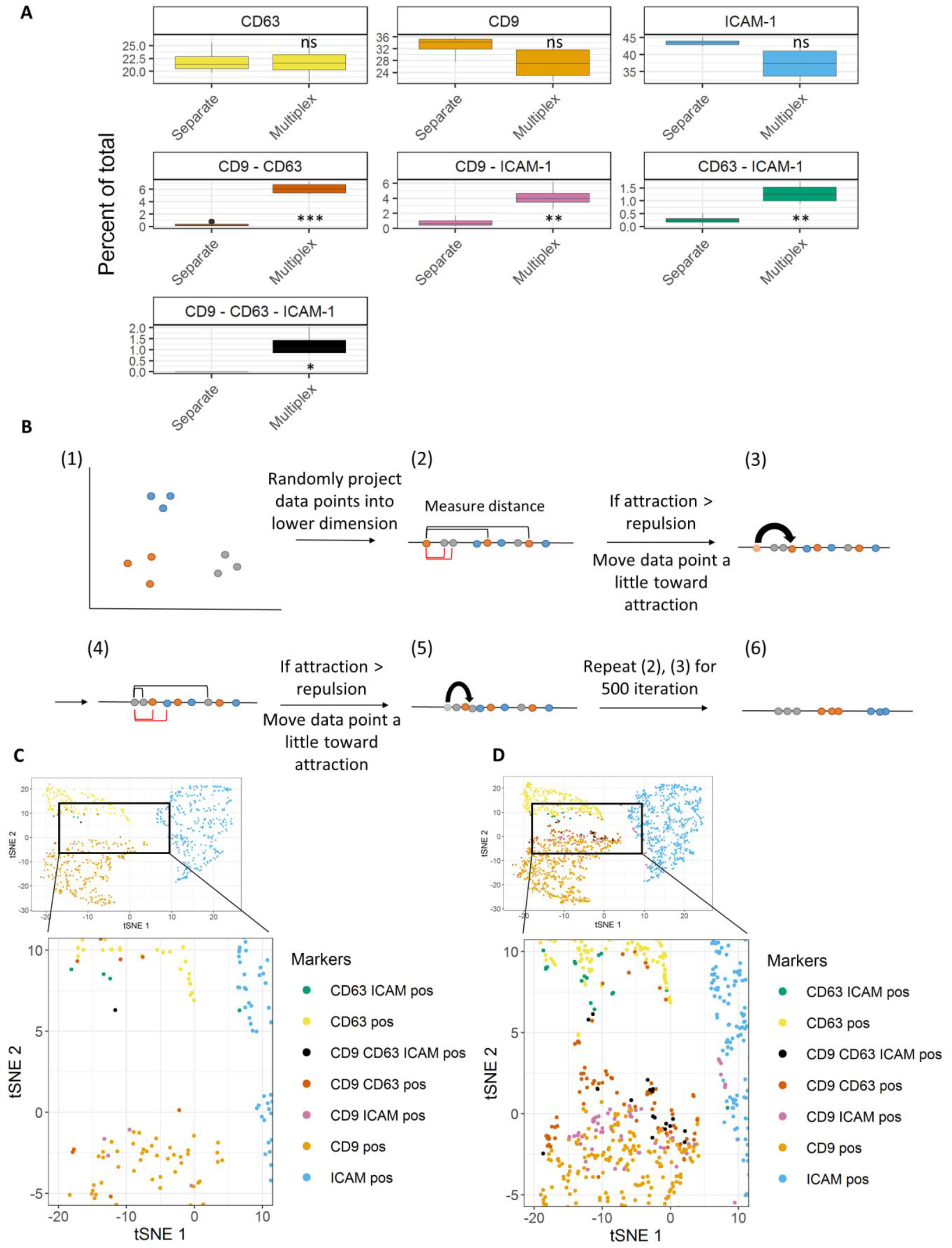
SBA applied for triple antibody labels, pEV were labeled using antibodies against Alexa 488 labeled anti-CD9, PE labeled anti-CD63 and Alexa 647 labeled anti-ICAM-1. Samples were either labeled separately and pooled or were labeled using the same antibodies in multiplex. A. Quantitative comparison of the separately labeled pEV sample with the multiplex labeled pEV sample, each label or label combination is presented as a percentage of the total detected bursts. B. Simplified schematic representation of how the t-SNE algorithm works. Here, 2D data will be reduced to 1D, while keeping the clustering intact. (1) Represents the dataset in 2D, clearly visualizing three distinct clusters. (2) Randomly project all data points into lower dimension. (3) Calculate the distance for each data point to all other data points. Data points close to each other in (1) attract, data points far from each other in (1) repel. (3) If attraction is larger than repulsion, move data point a little toward attractive points. In (4) and (5), steps (2) and (3) are repeated for the next data point. These steps are iterated through 500 times. (6) Shows the results of the dimensionality reduction maintaining the clusters of the original dataset (1). C. Visualizes the separately labeled sample data in a t-SNE plot, showing that mostly single labeled EV are present. The plot is zoomed in for detected multiple labels. D. Visualizes the multiplex labeled sample showing single and dual or triple labeled EV and zoomed in for the detected dual or triple labeled EV. (p-value < 0.05 considered significant.; ns, *, **, ***: significantly different from controls (not significant, *p* < 0.05, *p* < 0.01 and *p* < 0.001, respectively)).

**Figure 5.**
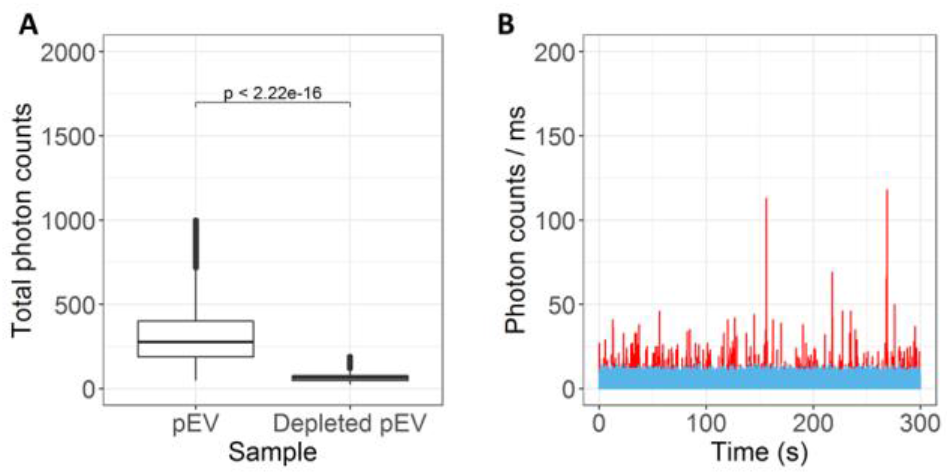
Three color SBA control experiments: CD9-CD81-CD63 (classic EV markers) depletion experiments showing a clear reduction of detected pEV signals. A. Comparison of pEV derived from plasma or from EV depleted plasma for three labels, showing a significant decrease for the total detected photons when depleting the plasma sample. B. Raw time traces measured using SBA, with in the *x*-axis time and in the *y*-axis counted photons of three channels. The blue trace represents the depleted sample, whereas the red trace represents the labeled pEV sample. *p* < 0.05 is considered as statistically significant as determined by student’s *t*-test

**Figure 6.**
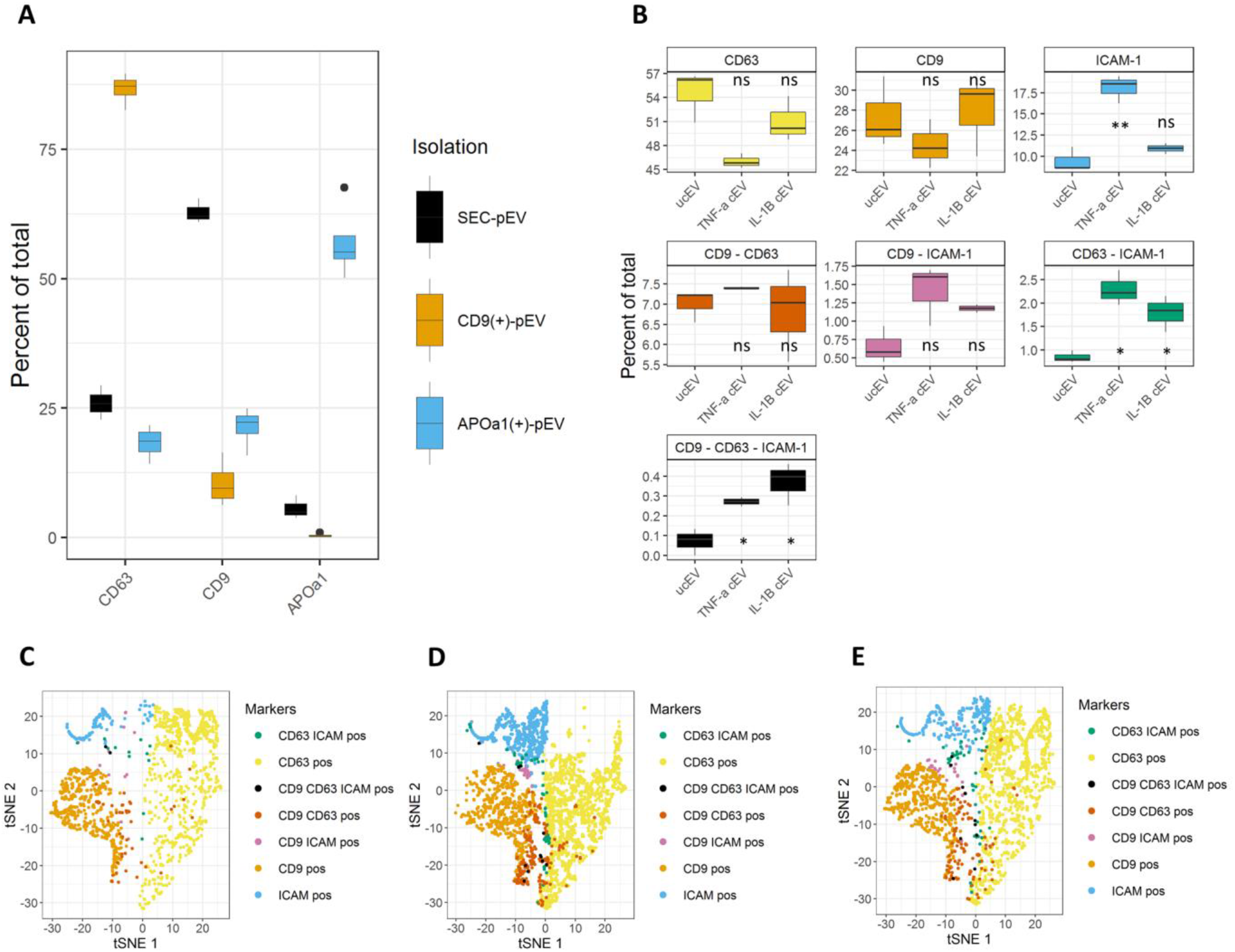
SBA applied as a fast quality control of a pEV preparation or for the detection of ICAM-1(+)EV subgroups derived from cell supernatant. **A**. Comparison of single step pEV SEC isolation (SEC-pEV) with ODG pEV isolation using SBA. CD9(+)-pEV, SEC isolated pEV from ODG fractions 9 and 10; APOa1(+)-pEV: SEC isolated APOa1 positive lipoproteins coming from ODG fractions one to three. (n = 3) **B**. SBA comparison of cEV isolated from cell conditioned media coming from HUVEC either untreated (ucEV) or treated with 10 ng/ml TNF-α (TNF-a cEV) or treated with 10 ng/ml IL-1β (IL-1β cEV) for 24 hours. (n = 3). **C-E** Visualization of t-SNE clusters for the different isolated cEV, with C. being the clusters defined for ucEV, D, for TNF-a cEV and E. for IL-1β cEV. One-way analysis of variance was used to compare ucEV with TNF-α cEV and IL-1β cEV. (*p-*value < 0.05 considered significant; ns, *, **: significantly different from controls (not significant, *p* < 0.05 and *p* < 0.01, respectively)).

The proposed approach allows the classification of EV sub-population, furthermore t-SNE clustering together with classification data allows the visualization of unique EV subtypes. This only requires a small sample size and allows for a quick and reproducible investigation of different EV subtypes.

### Depletion of classical EV markers confirms EV specificity of SBA signals

To further confirm that with the SBA approach EV-associated surface proteins rather than non-EV events are detected, we depleted EV from plasma by using magnetic beads conjugated with antibodies against three different transmembrane tetraspanin proteins CD9, CD63 and CD81 and then labeled the sample with three antibodies (figure 5).

Subsequently, SBA analysis indeed revealed a significant decrease (*p*<0.001) in the EV depleted fraction as compared to the bulk, confirming that with SBA only EV-associated (surface) markers are detected (figure 5A). These differences were also clearly visualized when comparing the raw data time traces of the depleted sample with a non-depleted sample (figure 5B). NTA also confirmed a depletion of 74% of the detected particles from crude fractions (SI figure 7A). Similar results were obtained for dual labeled samples (SI figure 7). Taken together, these results clearly indicate that the SBA-based approach can be used to detect one or several markers on a single EV allowing profiling of single EV into subgroups.

### Quality control of pEV and biomarker discovery using SBA

The capability of SBA based assays to perform EV quality control is demonstrated using different pEV fractions including a single step SEC isolated EV (SEC-pEV), still containing lipoproteins as already shown in the MISEV based quality control experiments and two highly purified EV fractions of ODG-SEC isolation procedures (APOa1(+)-pEV fractions and CD9(+)-pEV fractions). Figure 6 depicts the percentage of EV containing each marker as detected by SBA for the different EV preparations. Figure 6A shows the comparison of pEV preparations after a single step SEC procedure or a combination of SEC and ODG for three different markers CD63, CD9 and APOa1. Three different sample types were analyzed. The SEC-pEV sample results from a single SEC isolation, the APOa1(+)-pEV sample consists of ODG fractions 1-3 containing APOa1 (figure 1D), while the CD9(+)-pEV sample consists of the ODG fractions 9 and 10 containing CD9, as detected by western blot. CD63 was enriched for the CD9(+)-pEV fraction (86 ±4%), whereas it was comparable for the other two isolation methods (for SEC-pEV 26 ±3% and for APOa1(+)-pEV 18 ±4%). CD9 was mainly enriched in the SEC-pEV fraction (63 ±2%). The ODG fractions had lower percentages of CD9, namely 9 ±3%.and 21 ±5% for CD9(+)-pEV and APOa1(+)-pEV, respectively. APOa1 is clearly present if the one step SEC isolation procedure (6 ±2%) is applied (Figure 6A). Upon performing the ODG isolation step, it is depleted from the CD9(+) pEV fraction (0.44 ±0.40%) and enriched in the APOa1(+)-pEV fraction (58 ±9%). The CD9-CD63 dual labels ranged from 1 to 6%. These results show that SBA can perform fast and critical quality control measurements of EV isolates, using only a small sample aliquot.

In order to illustrate the applicability of the SBA assay in single EV biomarker research, we applied it to identify specific EV subpopulations associated with the induction of inflammation in cell culture. tumor necrosis factor alpha (TNF-α) and interleukin 1 beta (IL-1β) are both inflammatory cytokines and often used to simulate inflammation *in vitro*.^13^ We have previously shown that (TNF-α) treated human umbilical vein endothelial cells (HUVEC) produce EV carrying most of the adhesion markers as a hallmark of inflammation and making ICAM-1(+)-EV as a key candidate subpopulation of inflammation-associated EV.^14–16^ Here, using three fluorescently labeled antibodies (Alexa 488 labeled anti-CD9, PE labeled anti-CD63 and Alexa 647 labeled anti-ICAM-1), we applied our SBA assay to compare the inflammation-associated EV subgroup (ICAM-1(+)-EV) that is released by HUVEC triggered with two different types of inflammatory stimuli (IL-1β and TNF-α) (Figure 6B). As shown in figure 6B, the number of CD9(+)-EV released from HUVEC before (unconditioned (ucEV)) and after treatment (TNF-α or IL-1β cEV) with both stimuli was not significantly changed. In a similar manner, cells, undergoing IL-1β and TNF-α stimulation, release a similar number of CD63(+)EV as compared to untreated cells.

However, when looking at the inflammation associated EV subgroups carrying ICAM-1, we observed that the number of ICAM-1(+)-EV were significantly increased upon TNF-α stimulation, as compared to cells treated with IL1-β. Moreover, with our SBA approach the further classification of inflammation-associated EV subgroups based on multiple markers (two or three) was possible. Upon stimulation with either IL-1β or TNF-α, the number of CD63-ICAM-1 (+) EV were significantly increased as compared to ucEV. Interestingly, no significant differences were observed for the two other subgroups, *i.e*. CD9-CD63(+)-EV and (ii) CD9-ICAM-1(+)-EV. The number of EV carrying three markers (CD9-CD63-ICAM-1(+)-EV) were also increased after inflammatory stimulation as compare to ucEV.

The classified t-SNE defined clusters are visualized in figures 6C-E. Again, the ucEV samples shows a low presence of ICAM-1 and ICAM-1 detected subpopulations (figure 6C), whereas ICAM-1 and its subpopulations are clearly more abundant when triggered with the inflammatory stimuli TNF-α (figure 6D) or IL-1β (figure 6E).

Differences in detected ICAM-1 and ICAM-1 together with CD9 and/or CD63 labels on EV subgroups can be found based on the inflammatory stimulus (TNF-α or IL-1β) that was given to the same cells of origin, clearly illustrating that the production of different EV subsets is induced by different triggers. Overall these results clearly demonstrate the power of single EV profiling as well as EV subgroup visualization by applying our proposed SBA approach.

## Discussion and conclusion

EV are an important and easily accessible source of diagnostic information. However, EV characterization is still challenging since many of the available isolation methods have different effects on their quality. Furthermore, due to their nanosize, EV subgroup heterogeneity is masked by the conventionally used analysis methods on bulk EV samples. Several studies have shown the importance of detecting EV subpopulations and how these subpopulations may have a different cargo and/or different biological effects.^15–18^ However, the detection of these EV subpopulations is challenging. In addition, EV characterization is of crucial importance as many EV isolation procedures in use often co-isolate proteins and other biological entities, especially when applied to complex bio fluids, and as a consequence complicating EV biomarker discovery studies.^5,19,20^ To address these challenges, we applied the SBA approach for the high resolution detection of diagnostically relevant EV subpopulations.

The International Society for Extracellular Vesicles (ISEV) provides guidelines for standardized EV based experiments referred to as ‘minimal information for studies of extracellular vesicles’ (MISEV).^6^ Additionally, EV-track is an online platform furthering the standardization and increasing transparency of EV-reporting experiments.^21^ One of the most important criteria in these guidelines pertains to the quality control of isolated EV samples. Often the EV isolation methods depend on the source of the sample, the goal of the experiment as well as the amount of sample that is available. Despite the availability of a wide variety of EV isolation methods, the resulting EV preparations may be very different in achieved purity affecting their diagnostic potential. Indeed, Freitas *et al*. analyzed the presence of glycosylated EV populations using different isolation approaches and did show that ODG isolation as well as SEC methods provided enhanced EV glycoprotein yields as compared to other isolation methods.^22^ Similarly, Van Deun *et al*. compared four different EV isolation protocols and demonstrated that higher protein yields are not indicative of higher EV yields.^5^ In addition, they found that ODG purification of EV can efficiently remove known contaminating plasma proteins from EV preparations. The latter approach results in a pure EV fraction with a unique omics profile that is not found when applying other isolation methods.

Techniques available for the detection of single EV are limited, but are needed to improve the characterization and profiling of EV. Moreover, the currently used standard EV characterization techniques are time-consuming and only provide information on the bulk of EV isolates. Previous studies using fluorescence correlation spectroscopy and SBA have shown that it is possible to accurately characterize EV labeled with one labeled antibody as well as genetically engineered EV with green fluorescent protein (GFP) and mCherry.^9,23^ To our knowledge, SBA was not used for multidimensional characterization of single EV, nor for identifying unique EV subpopulations that are diagnostically relevant. The use of a small sample volume and straightforward, fast measurements and data analyses make SBA a powerful tool for EV characterization. Here, we propose an SBA approach that can be used two critical tools. It allows for a fast and reliable quality check of isolated EV samples, that can be used as a standard quality control test, thereby replacing time consuming and sample consuming standard EV characterization techniques. And even more important, we showed that the proposed SBA method can be applied to identify and quantify multiple parameters of single EV enabling the identification of unique and diagnostically relevant EV subpopulations at the single EV level.

### SBA enables sizing of single EV

SBA enables the sizing of both single 100 nm FITC labeled silica beads and single 100 nm TetraSpeck™ beads, resulting in a similar distribution to that when using NTA. Silica beads (n∼1.42) have a refractive index that is comparable to that of EV (n<1.42), showing the possibility of measuring the size of fluorescent nanobeads that share a physical characteristic with EV.^24^ Size characterization can be done using SBA. Our results are in correspondence with the measured size of the beads as well as with EV size and are in line with those previously demonstrated by Wyss *et al*..^9^

### SBA as a reliable quality control tool

The quality control of isolated EV samples aligns with previous studies. pEV isolated by using a single step SEC isolation were found to contain EV markers as well as APOa1-positive lipoproteins, as demonstrated by both western blot and TEM. The identification of these co-isolates is of importance as shown by Van Deun *et al*., reporting that these co-isolates are capable of interfering with the downstream analyses of EV and their associated biomarkers.^5^ Additionally, SBA was able to identify the presence of both CD9(+)-pEV and APOa1(+)-lipoproteins as a separate group. Apart from identifying these important co-isolates, SBA would also enable the detection of specific nanoparticle subgroups within a sample. ODG is suggested as a reliable method to separate EV and lipoproteins. The increased purity was confirmed by western blotting, identifying APOa1(+) fractions, indicative of lipoproteins, and CD9(+) fractions, indicating EV. This increased purity is further substantiated by TEM analysis of CD9(+)-pEV fractions and our results are in accordance with previous results reported by Van Deun *et al*. as well as others.^4,5,25^

Similarly, results obtained with SBA enabled the comparison of different isolation procedures for pEV. Comparison of these isolation procedures as a percentage of total detected bursts clearly shows that CD63 is enriched in the CD9(+)-pEV fractions, which is in line with the results reported by Van Deun *et al*..^5^ A decreased CD9 level in the CD9(+)-EV fractions was detected by SBA, but this can be explained by the additional SEC step that is required to exchange the viscous Optiprep™ buffer to PBS enabling diffusion-based measurements using SBA. This could reduce the yield of CD9 which is known to be more abundantly present on smaller EV. This is in accordance with Kowal *et al*., who showed that CD9 can be detected in both large and small EV but is predominantly present on small EV.^26^ Moreover, SBA confirmed the presence of APOa1 in the SEC-pEV fraction, its absence in the CD9(+)-pEV fraction and its enrichment in the APOa1(+)-pEV fractions. Hence SBA detects elegantly lipoprotein contaminations in patient plasma EV isolates More importantly, SBA results are collected in a fast (*i.e*. measurement possible within 60 seconds) and reliable way using only a small aliquot of the EV isolate and providing information on single EV, which is not feasible when using western blot or enzyme-linked immunosorbent assay (ELISA).^9^ To collect more data points we opted for measurements between 5 and 15 minutes, which is still significantly faster than the standard time for western blot or ELISA. SBA also enables for a reliable (digital) quantification of the bursts, that can easily be translated into a clinical setting where quality control on every EV sample is required.

### Multidimensional SBA as a tool to identify diagnostically relevant EV subpopulations

The possibility of detecting multiple labels on single EV was evaluated in different stages. First, TetraSpeck™ beads were used as a model to show that SBA can identify multiple labels on single particles, a clear difference compared to a single labeled sample was observed. Next, pEV were labeled with Alexa 488 labeled anti-CD9 and Alexa 647 labeled anti-ICAM-1 antibodies. The results clearly show that both single-labeled EV (CD9 or ICAM-1 (+)) EV can be distinguished, and that both labels are present on the EV surface. Additionally, triple labels were evaluated using a multiplex labeled and separately labeled and pooled experiment, where Alexa 488 labeled anti CD9, PE labeled anti-CD63 and Alexa 647 labeled anti-ICAM-1 were used as antibodies. With t-SNE we visualized different EV subgroups present in a sample. Three main single-labeled EV clusters were differentiated, together with multiple labeled EV. Quantitative analysis revealed that the multiple labeled EV accounted for 6% of the total detected EV. These low amounts of multiple labeled EV are similar to those observed in other studies.^27–30^ Again, our new SBA based approach was capable of specifically detecting multiple markers on a single EV allowing to sort single EV into different subgroups. In control experiments, using separately labeled and pooled EV samples a very low number of multiple labeled EV were observed using SBA. This is most likely due to the limitations of using the least squares method as an unmixing method, as a result of which some background signals are not accurately unmixed.^31^ However, significantly higher fractions of both dual labeled EV and triple-labeled EV were seen in the multiplex labeled EV fractions, which were additionally validated by applying an EV depletion step with anti-CD9, CD81 and CD63 antibodies, confirming that SBA specifically detected (multiple-)labeled EV. Fluorescence minus one controls are often used as both a negative control and a gating control in FACS experiments, enabling the quantification of the detector background.^32^ Also this approach confirmed the specificity of the observed SBA signals (SI Figure 8).

Current techniques usually analyze EV bulk fractions and are unable to capture the heterogeneity of specific EV subpopulations within these bulk EV samples. Identification of such specific EV subpopulations is important to identify novel diagnostically relevant EV subpopulations. As such we previously identified ICAM-1(+)-EV as an important EV subgroup that is released by inflammation-triggered endothelial cells and is potentially implicated in the development of cardiovascular diseases.^14,16^ Here, we demonstrated that the proposed SBA approach can be used to verify whether this important EV subset is significantly increased upon TNF-α stimulation. Interestingly, we further showed that the CD9-ICAM-1(+)-EV subgroup is not significantly increase upon stimulation, whereas the CD63-ICAM-1(+)-EV subgroup is, and that the detected triple labeled EV (CD9-CD63-ICAM-1(+)) are significantly increased upon inflammatory stimulation. Together, these findings clearly show that SBA can identify even low percentages of specific, diagnostically relevant EV subpopulations in an inflammatory context, which is not possible using current standard approaches.

### Further improvements for using SBA in single EV profiling

Our proposed SBA approach opens many opportunities in EV research field. We demonstrated that it can effectively be employed both as a fast quality control tool as well as a unique tool for profiling single EV. As such this approach has the potential to boost EV research in view of potential biomarker discovery. Subtle differences in EV composition are important for differences in effects in target cells. Nevertheless, some considerations need to be made to further improve the methodology. First, the choice of an appropriate fluorophore is of importance, preferably these fluorophores are bright and stable. In our experiments, we used antibodies labeled with Alexa labels or PE labels, which display excellent stability and brightness, allowing the detection well-defined bursts above the threshold.^27^ Also labeling of EV with fluorescent antibodies should be carefully designed, *e.g*. the binding efficiency of the specific antibody as well as the labeling efficiency of the antibodies themselves, *i.e*. what is the percentage of antibodies that are labeled. Moreover, appropriate smaller labeling molecules, such as single-chain antibodies, nanobodies or aptamers will increase the applicability of the SBA profiling of EV by decreasing possible steric interactions in all single EV techniques using labeled antibodies.^32-34^ Indeed, as reported by others, we also show that multiple labels are more likely to be found on larger EV as compared to smaller EV (SI figure 9A). This was also validated by adjusting the amount of labeling antibody, showing that 1.25 µg of antibody was the most optimal amount to detect EV with multiple labels (SI figure 9B). Correcting for spectral overlap procedures using the least squares method are often used in multiple applications like FACS. Nevertheless, it should be noted that these methods merely give an approximation of determining the correct signal for each channel.^31^

To improve and standardize multi-label detection on EV, suitable reference materials allow for a better evaluation of multiplex labeled EV phenotyping.^26^ Nevertheless, SBA measurements for single particles remain stable allowing for long measurements. The number of detected bursts remain consistent for at least an hour (SI figure 10A-E). SBA can be applied as a quantitative and reproducible technique. EV serial dilutions show a pronounced linear fit with an adjusted R^2^ of 0.99 (SI figure 10F). The proposed SBA methodology can be applied on labeled nanoparticles and considered as a stand-alone technique or as an orthogonal method supplementing other high-resolution technologies such as high-resolution FACS methodologies.

In conclusion, our results demonstrate that multidimensional SBA is a powerful tool that enables a fast and accurate evaluation of the purity of EV samples and allows for the multidimensional profiling of single EV. Thereby identifying unique and diagnostically relevant EV subpopulations. Therefore, applying the proposed SBA approach will significantly boost the diagnostic potential of EV in disease-related biomarker research.

## Materials and methods

### Antibodies and Reagents

Antibodies used for western blotting: CD9 (10626D, Invitrogen), CD63 (12-0639-42, Life technologies), CD81 (555675, BD biosciences), Annexin II (sc-28385, Sigma Aldrich), APOa1 (33505, Bioké), GM-130 (610822, BD biosciences) and rabbit anti-mouse HRP-conjugated secondary antibody (P026002-2, Dako). Antibodies used for SBA: Alexa 488 labeled mouse anti-human CD9 antibody (MCA469A488, Biorad), PE-labeled mouse anti human CD63 antibody (12-0639-42, Life technologies), Alexa 647 labeled mouse anti human ICAM-1 antibody (sc-107 AF 647, Santa Cruz) and Alexa 647 labeled mouse anti human APOa1 antibody (FAB36641R, Biotech). TNF-α (11343015) and IL-1β (11340013) were purchased from Immunotools GmbH. Sepharose CL-2B (17-0140-01 was purchased from VWR and Iodixanol (#07820) was purchased from Stemcell technologies.

### Cell culture

Human umbilical vein endothelial cells (HUVEC; BD Bioscience, cat # 354151) were seeded at passage 4 in a T75 flask with a density of 10.000 cells/cm^2^. Cells were grown in 15 mL EBM-2 (Lonza) supplemented with EGM-2 MV SingleQuot Kit (Lonza; except for the SingleQuot Kit fetal bovine serum) and 5% (V/V) FCS (LO CC-3202/6, Lonza) up to 70-75% confluency. Confluent cells were rinsed twice with PBS (Lonza) and treated with 10 ng/mL TNF-α (ImmunoTools GmbH, cat: 11343015) or Il-1β (Immunotools GmbH, cat: 11340013) in refreshed medium supplemented with 5% (V/V) exosome-depleted fetal bovine serum (A2720801, Gibco) for 24 h. Cells were incubated in a humidified atmosphere condition of 5% CO2 at 37 °C. Afterwards, supernatant was harvested after 24h from approximately 7.5 × 10^6^ total cells with more than ∼ 90% viability. Collected supernatant was centrifuged at 300 g for 10 min at 4 °C to remove cell debris. A second centrifugation step was done for 10 min at 2000 g at 4°C, to eliminate remaining debris and apoptotic bodies. Resulting supernatant was stored at −20 °C until further use.

### Isolation of fluorescently labeled EV from cell culture supernatant (cEV)

Before EV isolation, cell culture supernatant of HUVECs was concentrated to 1 mL using Amicon-Ultra 15 Centrifugal Filter Units (UFC901096, Merck) and incubated with 0.5 µg of Alexa 488 labeled anti-CD9, PE labeled anti-CD63 and/or Alexa 647 labeled anti-ICAM-1 antibodies. Then, fluorescently labeled EV were purified using sepharose CL-2B (17-0140-01, VWR) size exclusion chromatography (SEC) as described by Böing *et al*.^36^ One mL fractions were collected using an automatic fraction collector (Izon) and pooled EV-enriched fractions (F4 and F5) were used in further analysis.

### Isolation of fluorescently labeled EV from plasma (pEV)

Blood samples (50 mL) were collected in 9NC vacutainer tubes (367714, BD) from four healthy non-fasting adult volunteers in accordance with the relevant WHO guidelines and regulations. First 5 mL of drawn blood was discarded to avoid platelet activation. Platelet-free plasma was prepared by sequential centrifugation of citrated blood samples at 800 g for 10 min followed by 2 * 15 min at 2300 g, aliquoted and stored at −80 °C.

Citrated plasma was incubated with 1.25 µg of Alexa 488 labeled anti-CD9, PE labeled anti CD63 and/or anti APOa1 antibodies for 4h at RT while gently rotating. Labeled EV were isolated by SEC and Optiprep™ density gradient ultracentrifugation. SEC was performed as described above and the fractions 4, 5, 6 were collected for further purification using ODG as previously described.4 Briefly, different densities were prepared, using a working solution including 7 mL of Optiprep™ (#07820, Stemcell Technologies) with 1.4 mL of working buffer (0.25 M sucrose (200-334-9, VWR), 6 mM EDTA (E5134, Sigma Aldrich) and 60mM Tris-HCl (M151, VWR)). Next, the gradient solutions with concentrations of 5%, 10%, 20% and 40% were made by mixing the working solution and homogenization medium (HM) (0.25 M sucrose, 1 mM EDTA, 10mM Tris-HCl). Four mL of each density (3.5 mL for the 5% density) was stacked into an open polyallomer tube (337986, Beckman Coulter) using a Tecan freedom EVO, programmed with EVOware standard v1.4. The EV-containing fractions (F4, F5 and F6) were concentrated to 1 mL (UFC201024, Merck) before being pipetted on top of the ODG column. Afterwards, columns were spun at 100.000 g for 24 hours using a SW-28.1 rotor (k-factor: 276) in an optima XPN-80 ultracentrifuge (Beckman coulter), and subsequently fractions of 1 mL were collected. Finally, another SEC step was performed on the pools of different fractions (1-3 = APOa1(+)pEV, 9 and 10 = CD9(+)-EV) to exchange the buffer of EV fractions. In this step, pEV were isolated in a 0.32% citrate (S1804, Sigma-Aldrich) PBS buffer. One mL fractions were collected and fractions 4, 5, 6, 7 were pooled together and concentrated to 1mL (UFC201024, Merck) and stored at −80°C.

### pEV depletion using exosome isolation Pan kit

EV were depleted from citrated plasma using the exosome isolation kit pan human kit (130-110-912, Miltenyi Biotec) directed against a combination of CD9, CD63 and CD81 EV surface markers. Briefly, 50 µl isolation beads were incubated with 1 mL of citrated plasma for 1 h at room temperature. Next, pEV depleted and pEV enriched fractions were separated using a side-pull magnetic system.

### Nanoparticle tracking analysis

Concentration and size of the nanobeads (Silica and TetraSpeck™), cEV and pEV were analyzed using the NanoSight NS300 system (NanoSight, Malvern Ltd) equipped with a 532 nm laser. EV suspensions were injected into the sample chamber and measured three times for 60 s with a syringe flow rate of 80 au and the camera level and particle detection thresholds were adjusted at 14 and 9 in the software. Acquisitions were captured and analyzed using NTA software 3.2 (NanoSight. Malvern Ltd).

### Transmission electron microscopy

Size and morphology of EV and possible co-isolates within a sample were examined by TEM imaging. Sample preparation was adapted from Chen *et al*..^37^ Briefly, three droplets of the sample were placed on a clean Parafilm (291-1214, VWR), after which a Formvar Support Slot 2 × 1mm Ni Grid (FF2010-Ni, Electron Microscopy Sciences) was placed on top of the droplets and allowed to stand for 60 minutes to adsorb the fluid. The grid with adherent EV was washed 3x with PBS for 2 minutes and 5 times with Ultrapure water for 2 minutes. Droplets were fixed with 2% glutaraldehyde for 10 minutes, and then washed 5 times with Ultrapure water for 2 minutes. The grid was transferred to 2% uranyl acetate and allowed to stand for 15 minutes. The grid was then incubated in 0.13% methyl cellulose and 0.4% uranyl acetate for 10 minutes and dried at room temperature before examination with a Tecnai G2 Spirit BioTWIN (FEI, Eindhoven, The Netherlands). All solutions were filtered and UltraPure water was heated to release the CO2. Images were taken at 120 kV.

### Total protein quantification

The total protein concentration present in EV, plasma and cell lysates was quantified using a micro BCA™ protein assay kit (23235, ThermoFisher Scientific) following the manufacturer’s protocol. Optical density of standards and samples were measured at OD595 nm using a Multiskan™ FC Microplate Absorbance Reader (Thermo Scientific, Belgium). Plasma and cell lysate were prepared by mixing 100 µl citrated plasma with 100 µl RIPA buffer and 6 × 10^6^ HUVECs with 250 µl RIPA, respectively. Afterwards, lysates were centrifuged at 2000 g for 20 min and their supernatants were collected for western blotting analysis.

### Western blotting

One to five microgram total protein of cEV, pEV and plasma and cell lysates as positive controls were loaded on a 12% SDS-PAGE in loading buffer (50 mM Tris-HCl (pH 8.0)(M151, VWR), 150 mM NaCl (S7653, Sigma-Aldrich), 0.5% sodium deoxycholate (B20759, Alfa Aesar), 0.1% sodium dodecyl sulfate (M107, VWR), 1% NP-40 (74385, Sigma-Aldrich)) containing a protease inhibitor cocktail (05 892 970 001, Sigma-Aldrich). Electrophoresis was done at 200 V for 45 minutes. Next, the separated proteins were transferred onto a polyvinylidene fluoride membrane (Immobilon R, Merck Millipore Ltd) for 1 hour at 20 V. The membranes were blocked with 5% w/v fat free milk powder (Marvel) in PBS and were incubated overnight with 1:500 dilution of primary antibodies against conventional EV markers CD9 (10626D, Invitrogen), CD63 (12-0639-42, Life Technologies), CD81 (555675, BD Biosciences), Annexin II (sc-28385, Sigma-Aldrich), APOa1 (33505, Bioké) and GM-130 (610822, BD biosciences) at 4°C. After washing, membranes were incubated with rabbit anti-mouse horseradish peroxidase (HRP) conjugated antibody (P026002-2, Dako, 1:1000) as secondary antibody at RT for 1 h. Blots were developed using the WesternBright™ Quantum kit (K-12042-D10, Avansta) and visualized using an Amersham Imager 680 (GE Healthcare Life Sciences) system.

### Confocal microscopy measurements

Seven and a half microliter of either FITC labeled silica beads (100 nm, Si100-FC-1, NanoCS), TetraSpeck™ beads containing four well-seprarated excitation/emission peaks (360/430 nm, 505/515 nm, 560/580 nm and 660/680 nm) (100 nm, T7284, Life Technologies) were used in a 1/10.000 dilution or fluorescently labeled EV (10^8^ particles) were placed between two #1 coverslips (15737592, Thermofisher) using a spacer of 120 µm (654004, Sigma-Aldrich).

All SBA experiments were performed on a Zeiss LSM 880 confocal laser scanning microscope (Carl Zeiss, Jena, Germany) using a Zeiss C-Apochromat 63x/1.2 W Korr Objective with Milli Q water in FCS mode recording raw photon data. A 488/543/633 mirror beam splitter (MBS) was used in combination with an Argon-ion laser (488 nm, 4% power, 27 µW), a HeNe laser (543 nm, 7% power, 45 µW) and a HeNe laser (633 nm, 5% power, 80 µW). All experiments were done at room temperature.

To obtain the physical parameters of the focal volume, fluorescence correlation spectroscopy was carried out using a mixture of Atto488 and Atto655, with D = 400 µm^2^/s as a fixed diffusion coefficient, yielding the ω_r_ and ω_z_ calibration fraction.^38^

All SBA experiments consisted of at least 5-15 acquisitions of 60 seconds. These were recorded with a sampling time of 15 MHz. The Zen black software 2.3 (Zeiss) was used to set up and record all SBA measurements.

### Data extraction from raw FCS files

All high time-resolution photon data (Zeiss .raw format) was saved in the Zen black software 2.3. A burst search was performed on the sum of all photon channels (APBS-2c-noMFD or APBS-3c-MFD) after opening the data in the open source pulsed interleaved excitation analysis in the MATLAB (PAM) package (Muenchen).^39^ Extracting the burst info was done using burst search in the burst analysis tab of the PAM software. Sliding time window was selected as the smoothing method and the all photon burst search for two without multi-fluorescence detection ‘APBS 2c-noMFD’ or for three colors ‘APBS 3c-MFD’ was selected as burst search method. The parameters used in the burst search are: minimum photons per burst = 5, time window (µs) = 500, photons per time window = 5. After the burst search, data was opened in BurstBrowser (PAM) and all the bursts were exported as comma separated values into csv files.

### Data analysis – Single burst extraction and visualizations

Further data analysis starts by loading the csv file in R 4.02. (R core team (2013)) To quantify the cross-talk between channels, the least squares method was used with single labeled EV on the basis of equation (1).^40^

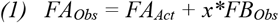

with *FA*_*Obs*_ the observed fluorescence in channel A, *FA*_*Act*_ the actual fluorescence in channel A and *FB*_*Obs*_ the observed fluorescence in channel B; *FA*_*Act*_ and the multiplication factor x are determined by the least squares approach.

After unmixing, all values smaller than 1 were put to equal 1. All values higher than 2000 counted photons (determined as outliers) in each channel were discarded. Thresholds were calculated by taking the mean of all the counted photons per channel and adding three times the standard deviation. After thresholding, data were classified for their respective fluorescent label using the calculated thresholds and plots were made for visualization and interpretation of the data.

The path of a detected molecule through the focal volume is related to the size of the molecule. Size calculations can be performed for the burst duration of each accepted individual burst using following equations, as was done by Wyss *et al*..^8^ The average diffusion time *τ*_*D*_ is given by:

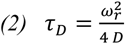

with *ω*_*r*_ the calibration factor of the focal volume in the plane perpendicular to the optical axis of the microscope and *D* the diffusion constant,

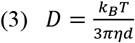

with *k*_*B*_ the Boltzmann constant, *T* the absolute temperature, *η* the viscosity of water and d the hydrodynamic diameter. The particle diameter *d* is experimentally determined as follows

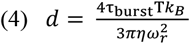

with *τ*_*burst*_ being the burst duration of a single defined burst. This burst duration is dependent on the path of the molecule through the focal volume, as this path is random, this can be employed for the sizing of the average nanosized particle.

In these equations *τ*_*D*_ takes into account the average diffusion time of all the detected photons in the time trace, whereas *τ*_*burst*_ will take into account the burst duration of a single defined burst.

A basic analysis R code that enables the data analysis and processing will be made available using the following link: https://github.com/Aptamer1/EV_SBA.

### Statistical Analysis

Data comparisons are presented as a mean ± SD of at least three independent experiments. Either student’s *t*-test was applied for the comparison between two groups or one-way analysis of variance (ANOVA) was applied for the comparison between multiple groups using R 4.02 to evaluate the statistical significance between different samples. Tests at value of *p* < 0.05 and were considered as statistically significant. NS represented as not significant, *p* > 0.05.

The visualization and detection of patterns within the data was done using t – distributed stochastic neighbor embedding (t-SNE).^10^ The Rtsne package was used to visualize local clusters.^41^ First the perplexity hyperparameter was optimized using a perplexity of 2, 5, 30, 50 and 100, to visualize the clusters optimally. For these data, a perplexity of 50, 500 iterations and a learning rate of 200 were chosen as optimal parameters allowing the differentiation of different nanoparticle populations. This algorithm allows the identification of detected EV subpopulations with relation to their fluorescent label and/or the average size. After performing t-SNE, data were labeled using the threshold classification.

### EV track

All relevant data of our experiments were submitted to the EV-TRACK knowledgebase (EV-TRACK ID: EV200010).^21^

## ASSOCIATED CONTENT

### Supporting Information

All figures can be found in the provided pdf file. SI figure 1: Low magnification TEM images

SI figure 2: Autocorrelation function to fir for τ_D_

SI figure 3: Visualization of triple labeled burst data obtained from BurstBrowser (PAM).

SI figure 4: Overview of t-SNE defined clusters

SI figure 5: t-SNE hyperparameter optimization for a triple labeled pEV sample

SI figure 6: t-SNE hyperparameter optimization for TetraSpeck™ beads data

SI figure 7: CD9 – CD81 – CD63 depletion experiments SI figure 8: Fluorescence minus one controls

SI figure 9: Detected EV with multiple labels

SI figure 10: Stability of a measurement of one hour and dilution series

## AUTHOR INFORMATION

### Author Contributions

S.K. and N.S. conceived the idea. S.K. performed the EV experiments. I.P. and J-P.T. performed the TEM experiments. S.K. did the SBA experiments and data analysis under supervision of J.H. and M.A.. S.K., M.A., J.H., L.M., B.H. interpreted the results. All authors took part in the discussion and writing.

## ACKNOWLEDGMENT

The authors thank Véronique Vastmans and Iris Reniers for the technical assistance.

---

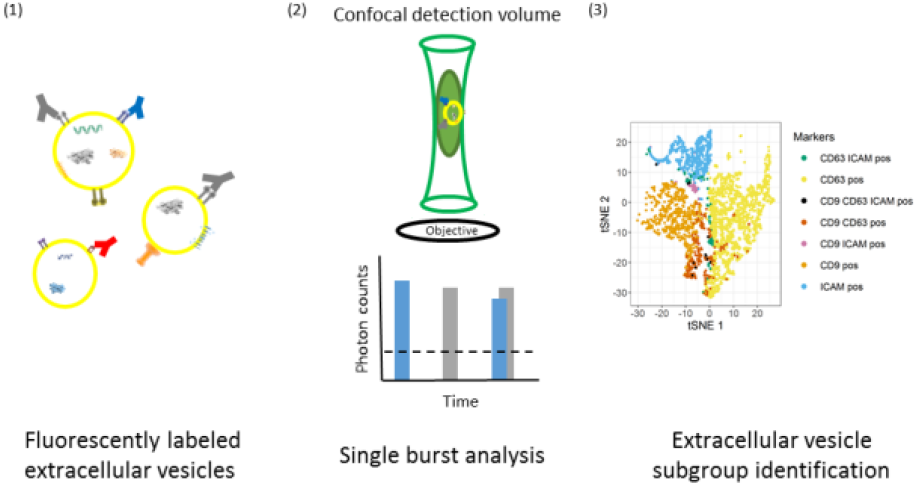

## REFERENCES

1. Raposo, G. & Stoorvogel, W. Extracellular vesicles: exosomes, microvesicles, and friends. J. Cell Biol. 200, 373–83 (2013).

2. Théry, C., Ostrowski, M. & Segura, E. Membrane vesicles as conveyors of immune responses. Nature Reviews Immunology 9, 581–593 (Nature Publishing Group, 2009).

3. Armstrong, D. & Wildman, D. E. Extracellular vesicles and the promise of continuous liquid biopsies. J. Pathol. Transl. Med. 52, 1–8 (2018).

4. Ramirez, M. I. et al. Technical challenges of working with extracellular vesicles. Nanoscale 10, 881–906 (2018).

5. Van Deun, J. et al. The impact of disparate isolation methods for extracellular vesicles on downstream RNA profiling. J. Extracell. vesicles 3, (2014).

6. Théry, C. et al. Minimal information for studies of extracellular vesicles 2018 (MISEV2018): a position statement of the International Society for Extracellular Vesicles and update of the MISEV2014 guidelines. J. Extracell. Vesicles 8, 1535750 (2019).

7. Chiang, C.-Y. & Chen, C. Toward characterizing extracellular vesicles at a single-particle level. J. Biomed. Sci. 26, 9 (2019).

8. Zander, C. et al. Detection and characterization of single molecules in aqueous solution. Appl. Phys. B Lasers Opt. 63, 517–523 (1996).

9. Wyss, R. et al. Molecular and Dimensional Profiling of Highly Purified Extracellular Vesicles by Fluorescence Fluctuation Spectroscopy. Anal. Chem. 86, 7229–7233 (2014).

10. Van Der Maaten, L. & Hinton, G. Visualizing Data using t-SNE. Journal of Machine Learning Research 9, (2008).

11. Andrews, T. S. & Hemberg, M. Identifying cell populations with scRNASeq. Molecular Aspects of Medicine 59, 114–122 (2018).

12. Acuff, N. V. & Linden, J. Using Visualization of t -Distributed Stochastic Neighbor Embedding To Identify Immune Cell Subsets in Mouse Tumors. J. Immunol. 198, 4539–4546 (2017).

13. Zhang, F. et al. IL-1/TNF-α inflammatory and anti-inflammatory synchronization affects gingival stem/progenitor cells’ regenerative attributes. Stem Cells Int. 2017, (2017).

14. Hosseinkhani, B. et al. Direct detection of nano-scale extracellular vesicles derived from inflammation-triggered endothelial cells using surface plasmon resonance. Nanomedicine Nanotechnology, Biol. Med. 13, 1663–1671 (2017).

15. Hosseinkhani, B., Kuypers, S., van den Akker, N. M. S., Molin, D. G. M. & Michiels, L. Extracellular Vesicles Work as a Functional Inflammatory Mediator Between Vascular Endothelial Cells and Immune Cells. Front. Immunol. 9, 1789 (2018).

16. Hosseinkhani, B., van den Akker, N. M. S., Molin, D. G. M. & Michiels, L. (Sub)populations of extracellular vesicles released by TNF-α –triggered human endothelial cells promote vascular inflammation and monocyte migration. J. Extracell. Vesicles 9, 1801153 (2020).

17. Tulkens, J., De Wever, O. & Hendrix, A. Analyzing bacterial extracellular vesicles in human body fluids by orthogonal biophysical separation and biochemical characterization. Nat. Protoc. 15, 40–67 (2020).

18. Vagner, T. et al. Large extracellular vesicles carry most of the tumour DNA circulating in prostate cancer patient plasma. J. Extracell. Vesicles 7, 1505403 (2018).

19. Dhondt, B. et al. Purification of urinary extracellular vesicles for uro-oncological biomarker studies using an iodixanol (Optiprep™) density gradient. Eur. Urol. Suppl. 16, e1078–e1079 (2017).

20. Dhondt, B. et al. Unravelling the proteomic landscape of extracellular vesicles in prostate cancer by density-based fractionation of urine. J. Extracell. Vesicles 9, 1736935 (2020).

21. Van Deun, J. et al. EV-TRACK: transparent reporting and centralizing knowledge in extracellular vesicle research. Nat. Methods 14, 228–232 (2017).

22. Freitas, D. et al. Different isolation approaches lead to diverse glycosylated extracellular vesicle populations. J. Extracell. vesicles 8, 1621131 (2019).

23. Corso, G. et al. Systematic characterization of extracellular vesicle sorting domains and quantification at the single molecule – single vesicle level by fluorescence correlation spectroscopy and single particle imaging. J. Extracell. Vesicles 8, 1663043 (2019).

24. van der Pol, E., Coumans, F. A. W., Sturk, A., Nieuwland, R. & van Leeuwen, T. G. Refractive Index Determination of Nanoparticles in Suspension Using Nanoparticle Tracking Analysis. Nano Lett. 14, 6195–6201 (2014).

25. Tauro, B. J. et al. Comparison of ultracentrifugation, density gradient separation, and immunoaffinity capture methods for isolating human colon cancer cell line LIM1863-derived exosomes. Methods 56, 293–304 (2012).

26. Kowal, J. et al. Proteomic comparison defines novel markers to characterize heterogeneous populations of extracellular vesicle subtypes. Proc. Natl. Acad. Sci. U. S. A. 113, E968–E977 (2016).

27. Morales-Kastresana, A. et al. Labeling Extracellular Vesicles for Nanoscale Flow Cytometry. Sci. Rep. 7, 1878 (2017).

28. Ricklefs, F. L. et al. Imaging flow cytometry facilitates multiparametric characterization of extracellular vesicles in malignant brain tumours. J. Extracell. Vesicles 8, 1588555 (2019).

29. Johnson, S., Banyard, A., Smith, C., Mironov, A. & McCabe, M. Large extracellular vesicles can be characterised by multiplex labelling using imaging flow cytometry. bioRxiv 2020 (2020). doi:10.1101/2020.02.07.938779

30. Burbidge, K. et al. Cargo and cell-specific differences in extracellular vesicle populations identified by multiplexed immunofluorescent analysis. J. Extracell. Vesicles 9, 1789326 (2020).

31. Li, H. C. & Chang, C. I. Linear spectral unmixing using least squares error, orthogonal projection and simplex volume for hyperspectral images. in Workshop on Hyperspectral Image and Signal Processing, Evolution in Remote Sensing 2015-June, (IEEE Computer Society, 2015).

32. Tung, J. W. et al. Modern Flow Cytometry: A Practical Approach. Clinics in Laboratory Medicine 27, 453–468 (2007).

33. Chuo, S. T. Y., Chien, J. C. Y. & Lai, C. P. K. Imaging extracellular vesicles: Current and emerging methods. Journal of Biomedical Science 25, (2018).

34. Tian, Y. et al. Protein Profiling and Sizing of Extracellular Vesicles from Colorectal Cancer Patients via Flow Cytometry. ACS Nano 12, 671–680 (2018).

35. Mastoridis, S. et al. Multiparametric analysis of circulating exosomes and other small extracellular vesicles by advanced imaging flow cytometry. Front. Immunol. 9, 1583 (2018).

36. Böing, A. N. et al. Single-step isolation of extracellular vesicles by size-exclusion chromatography. J. Extracell. Vesicles 3, (2014).

37. Chen, C. L. et al. Comparative and targeted proteomic analyses of urinary microparticles from bladder cancer and hernia patients. J. Proteome Res. 11, 5611–5629 (2012).

38. Kapusta, P. Absolute Diffusion Coefficients: Compilation of Reference Data for FCS Calibration. (2010). doi:10.1209/0295-5075/83/46001

39. Schrimpf, W., Barth, A., Hendrix, J. & Lamb, D. C. PAM: A Framework for Integrated Analysis of Imaging, Single-Molecule, and Ensemble Fluorescence Data. Biophys. J. 114, 1518–1528 (2018).

40. Nolan, J. P. & Condello, D. Spectral flow cytometry. Curr. Protoc. Cytom. CHAPTER, Unit1.27 (2013).

41. Krijthe, J. H. {Rtsne}: T-Distributed Stochastic Neighbor Embedding using Barnes-Hut Implementation. (2015).

