## Supplemental information for "Machine learning-based clustering of nanosized fluorescent extracellular vesicles"

### Supporting information

Machine learning-based clustering of nanosized fluorescent extra-cellular vesicles

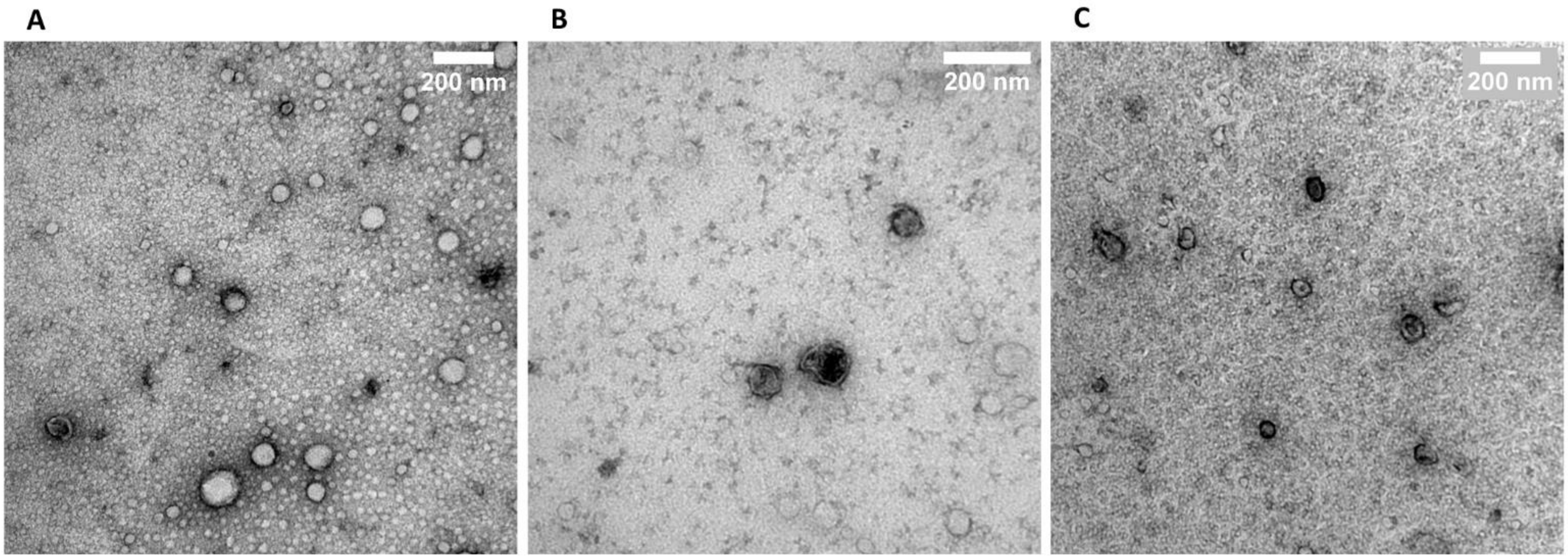


SI figure 1: Low magnification TEM images showing the sample heterogeneity. A. SEC-pEV visualization, showing a clear presence of co-isolated contaminants within the sample (Scale bar 200 nm). B. CD9(+)-pEV fraction visualizing a clear decrease of the background and contaminants in the pEV sample as well as the presence of pEV (Scale bar 200 nm). C. cEV clearly showing the presence of cEV as well as a low background and low presence of co-isolates (Scale bar 200 nm).

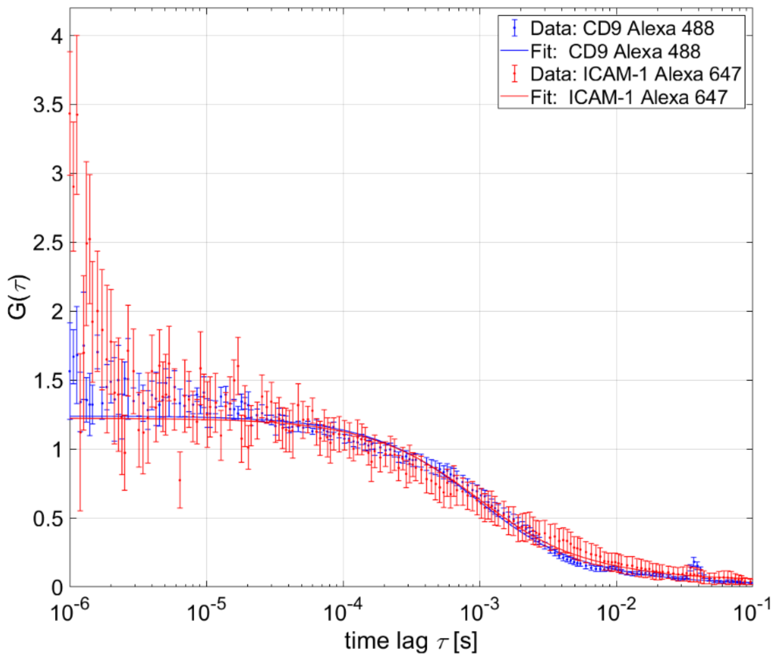


SI figure 2: Fluorescence correlation spectroscopy time traces analyzed for channel 488 (anti-CD9 Alexa 488) and channel 647 (anti-ICAM-1 Alexa 647). The autocorrelation function is fit with the FCS_TauD fit model in FCSFit (PAM). A global $\tau_D$ of 1.75 ms was found, corresponding with a mean size of 85 nm.

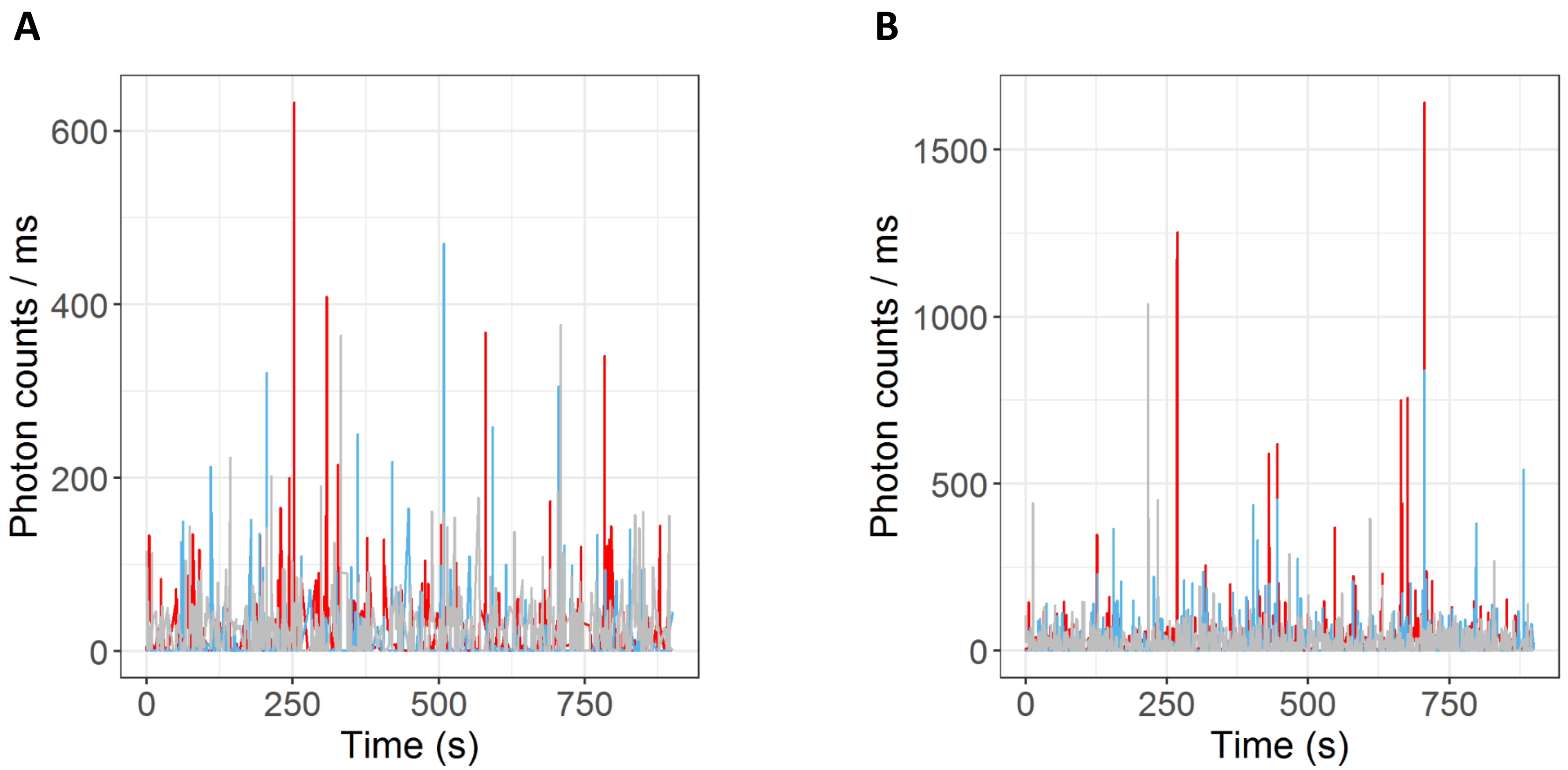


SI figure 3: Visualization of triple labeled burst data obtained from BurstBrowser (PAM). A. Represents the separately labeled and pooled pEV sample and B. represents the triple labeled pEV sample. Anti-CD9 Alexa 488 (red), anti-CD63 PE (blue) and anti-ICAM-1 Alexa 647 (gray) were used as antibodies. This data is further analyzed by classifying the bursts and clustering with t-SNE.

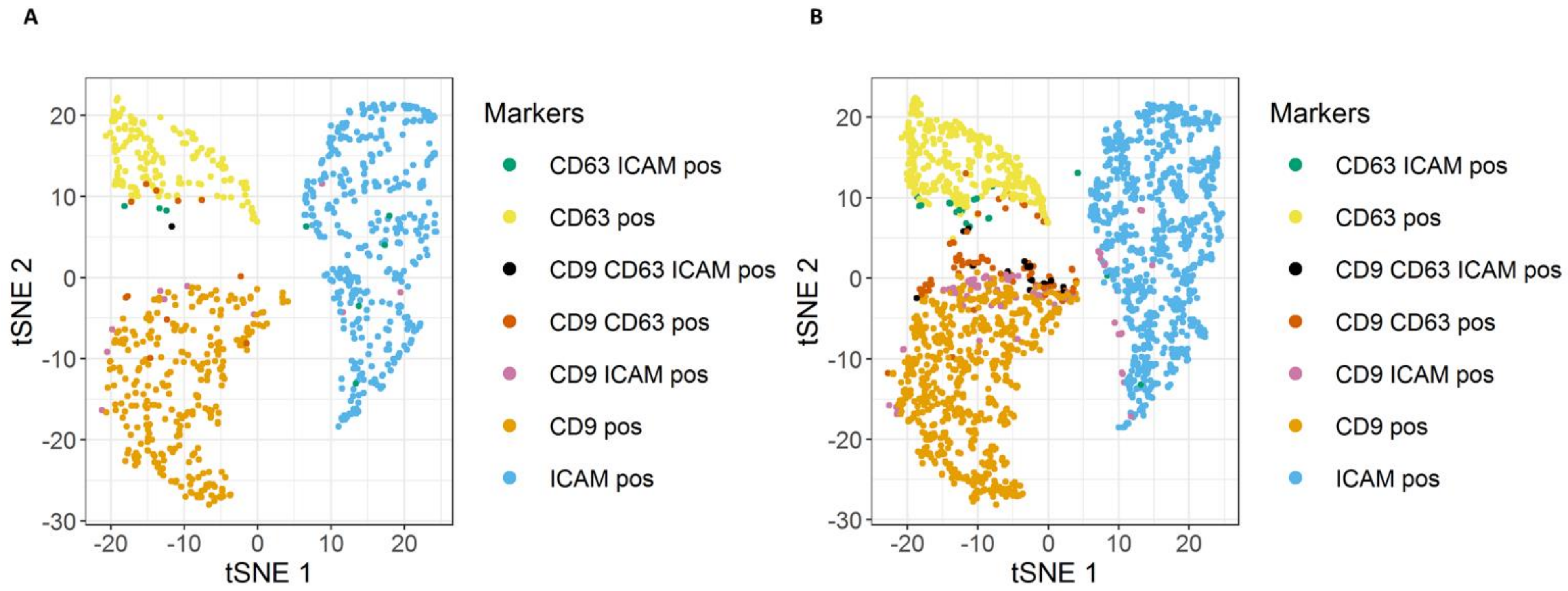


SI figure 4: Overview of the t-SNE defined clusters, with A. the separately labeled and pooled EV sample and B. the triple labeled (multiplex) EV sample.

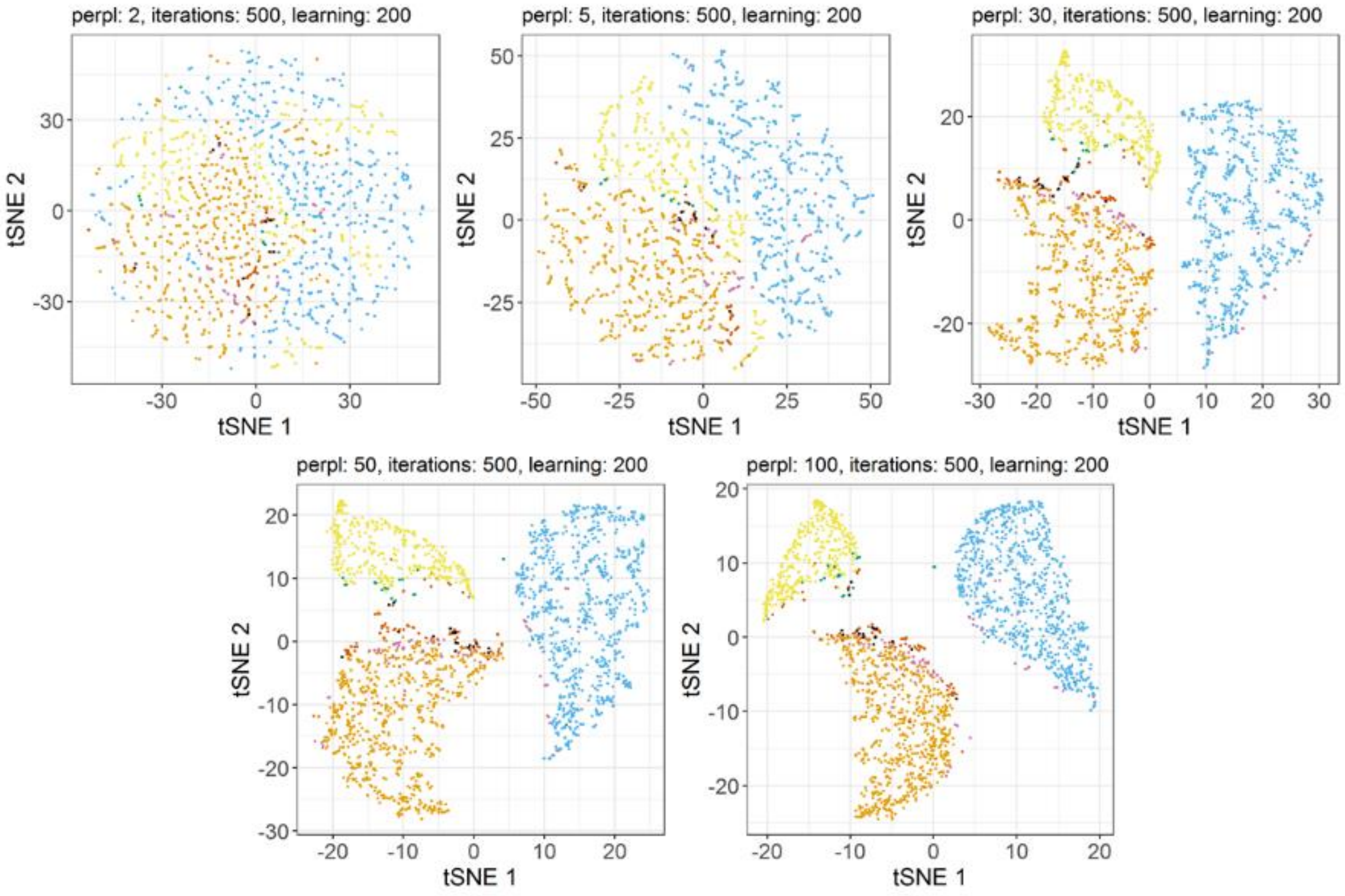


SI figure 5: t-SNE is a dimensionality reduction algorithm which allows to find structures or patterns within high dimensional data. Optimization of the t-SNE perplexity parameter was done using an automated script for a triple labeled pEV sample. Perplexity is an important optimizable hyperparameter of t-SNE and affects how well the data is clustered. Each dot within the plots represents a dimensionality reduction of the size of the individual EV as well as their detected signal in each of the three channels. The figure visualizes perplexities ranging from 2 to 100 while keeping the amount of iterations and the learning rate constant. Lower perplexity numbers are not able to separate the clusters within the detected pEV sample at all, whereas higher perplexities separate them almost completely. Here, a perplexity of 30 or 50 separates the main clusters within the data quite well. For our approach a perplexity of 50 was chosen.

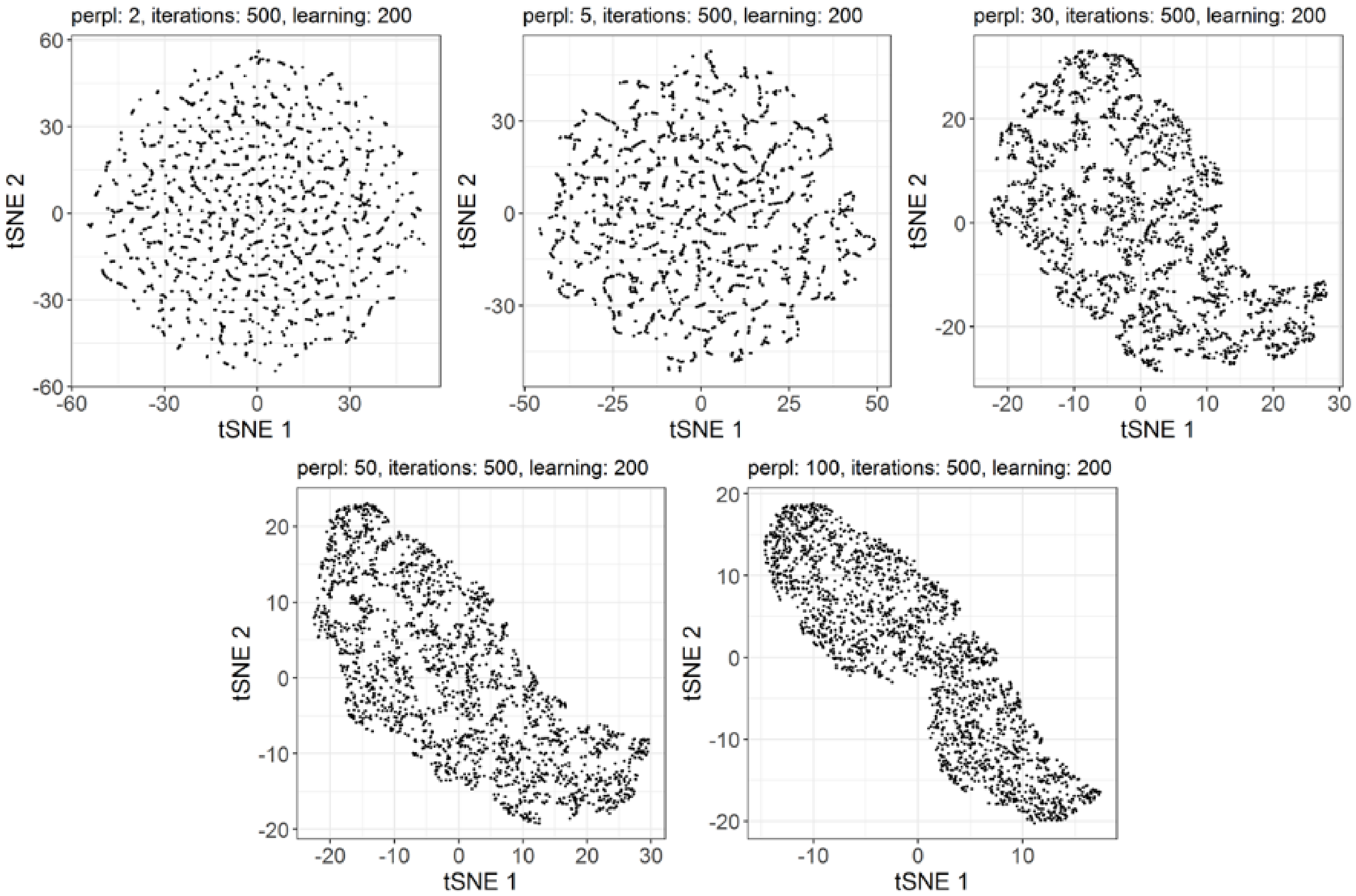


SI figure 6: Visualization of t-SNE applied for measured TetraSpeck$^{TM}$ beads, with 500 iterations, a learning rate of 200 and perplexities ranging from (2-100), similar to the EV data. These graphs clearly show that the lower perplexities are not capable of finding any clusters within the TetraSpeck$^{TM}$ data. A perplexity of 100 did show a tendency to a potential pattern within the data, however no real pattern separation into subgroups was achieved. This tendency to a pattern is most likely triggered by differences in intensities for these beads.

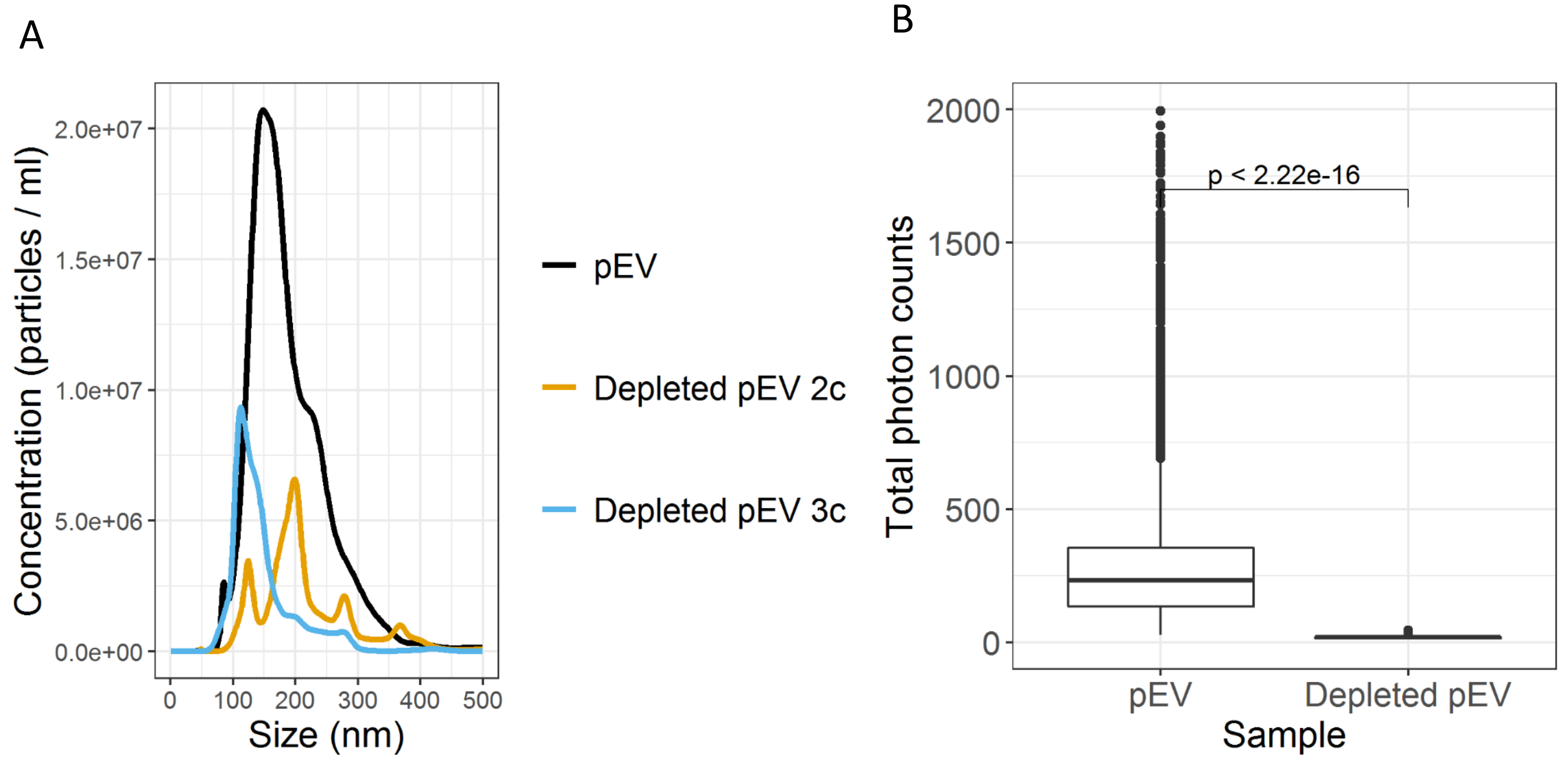


SI figure 7: CD9 – CD81 – CD63 depletion experiments showing a clear reduction of detected pEV. A. NTA results of pEV compared to the two depleted pEV samples, with pEV having a concentration of 2.28E+09 particles/mL, the depleted pEV 2c a concentration of 5.82E+08 particles/mL and the depleted pEV 3c a concentration of 5.98E+08 particles/mL. B. Comparison of dual labeled pEV derived from plasma or from EV depleted plasma for two labels, showing a significant decrease of total detected photons upon depletion.

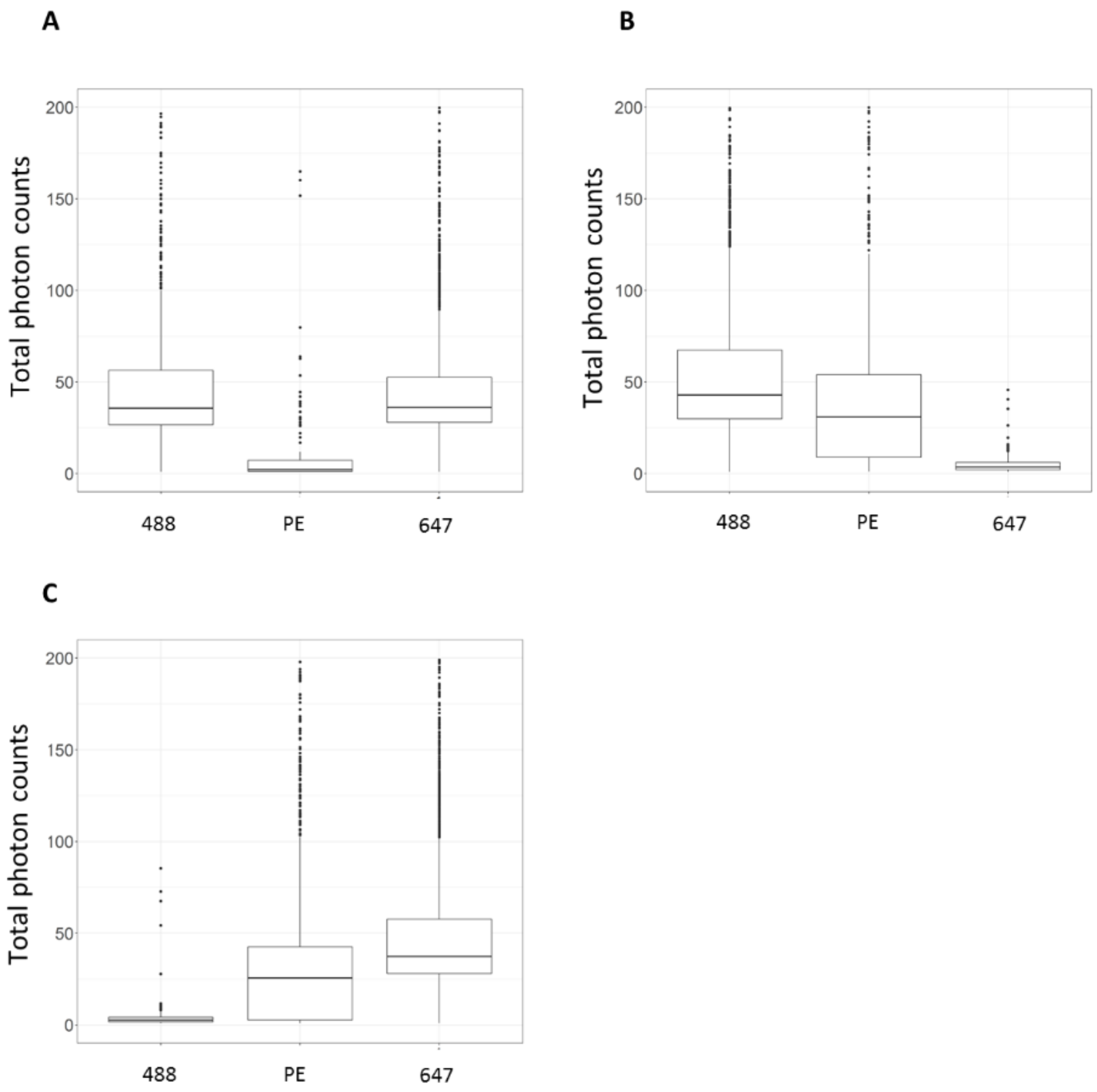


SI figure 8: Fluorescence minus one controls showing the amount of counted photons in each channel. Each experiment is recorded in three channels, while the sample is only labeled with a combination of only two labeled antibodies (Alexa 488 labeled anti-CD9 = 488, PE labeled anti-CD63 = PE, Alexa 647 labeled anti-ICAM-1 = 647). A. Visualizes the signal of a pEV sample labeled with A488 and A647, while the signal of the PE channel is decreased. B. Demonstrates the signal of a pEV sample labeled with A488 and PE, while the signal of the A647 channel is decreased. C. Showing the signal of a pEV sample labeled with PE and A647, while the signal of the A488 channel is decreased.

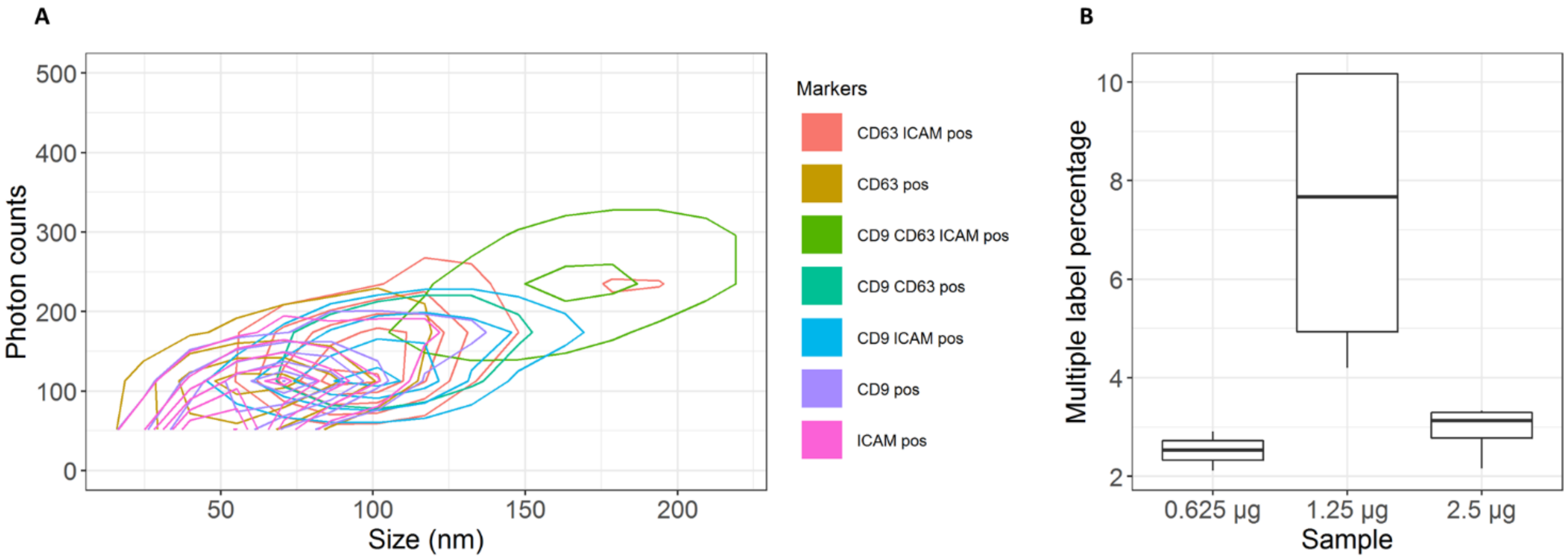


SI figure 9: Detected multi-label EV have an increased size and is affected by the amount of antibodies used. A. 2D-kernel density plot, visualizing the probability of detecting a marker within the data, with in the x-axis the size (nm) and in the y-axis the photon counts of three channels. This plot shows that there is a correlation between the size and the amount of photon counts (Pearson correlation of 0,83). Additionally, the lowest EV sizes are mainly labeled with a single antibody, whereas bigger EV sizes are more likely to have multiple labels detected. It is likely that this phenomenom can be described by sterical hindrance, as these smaller EV are around the size of 80 nm and the larger EV are around 130 nm. B. Antibody titration experiments using different antibody amounts, namely 0.625 µg,1.25 µg or 2.5 µg of antibodies. All multiple labels are quantified as a percentage of the total. There is a signficant difference when comparing 0.625 µg with 1.25 µg (*p*-value = 0.012) and a signficant different when comparing 2.5 µg with 1.25 µg (*p*-value = 0.02). It is likely that 0.625 µg does not have enough antibodies to label all multiple labels, whereas 2.5 µg effects can be explained by a higher background, different antibody affinities as well as sterical hindrance effects. A $p$-value $< 0.05$ is considered as statistically significant as determined by one way analysis of variance.

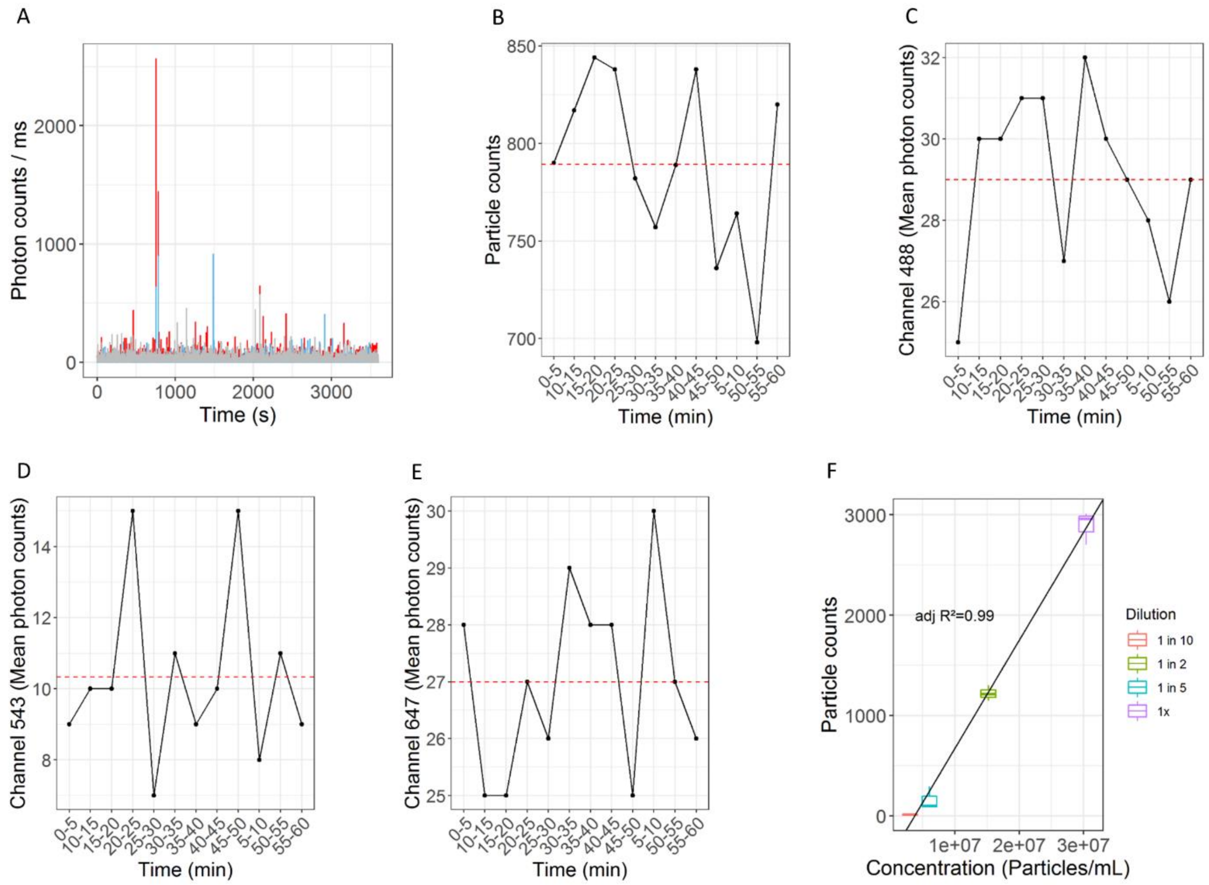


SI figure 10: Stability and linearity of detected pEV. A. Fluorescent pEV fluctuations measured during a time period of one hour. B. The number of detected particle counts every 5 minutes during a measurement period of one hour, the red dashed line visualizes the mean. C,D,E. Mean detected photons of each channel every 5 minutes during a measurement of one hour, the red dashed line visualizes the mean. It is clear that the number of detected bursts remains stable throughout the entire period. F. Dilution series of pEV measurements in a concentration range of 3E+07 – 3E+06 particles/mL as determined by NTA, showing a linear relationship with the burst analysis measurement with an adjusted $R^2$ of 0.99.
